# Non-Coding Genetic Analysis Implicates Interleukin 18 Receptor Accessory Protein 3′UTR in Amyotrophic Lateral Sclerosis

**DOI:** 10.1101/2021.06.03.446863

**Authors:** Chen Eitan, Aviad Siany, Elad Barkan, Tsviya Olender, Kristel R. van Eijk, Matthieu Moisse, Sali M. K. Farhan, Yehuda M. Danino, Eran Yanowski, Hagai Marmor-Kollet, Natalia Rivkin, Nancy Yacovzada, Shu-Ting Hung, Johnathan Cooper-Knock, Chien-Hsiung Yu, Cynthia Louis, Seth L. Masters, Kevin P. Kenna, Rick A. A. van der Spek, William Sproviero, Ahmad Al Khleifat, Alfredo Iacoangeli, Aleksey Shatunov, Ashley R. Jones, Yael Elbaz-Alon, Yahel Cohen, Elik Chapnik, Daphna Rothschild, Omer Weissbrod, Gilad Beck, Elena Ainbinder, Shifra Ben-Dor, Sebastian Werneburg, Dorothy P. Schafer, Robert H. Brown, Pamela J. Shaw, Philip Van Damme, Leonard H. van den Berg, Hemali P. Phatnani, Eran Segal, Justin K. Ichida, Ammar Al-Chalabi, Jan H. Veldink, Project MinE ALS Sequencing Consortium, NYGC ALS Consortium, Eran Hornstein

## Abstract

The non-coding genome is substantially larger than the protein-coding genome but is largely unexplored by genetic association studies. Here, we performed region-based burden analysis of >25,000 variants in untranslated regions of 6,139 amyotrophic lateral sclerosis (ALS) whole-genomes and 70,403 non-ALS controls. We identified Interleukin-18 Receptor Accessory Protein (IL18RAP) 3′UTR variants significantly enriched in non-ALS genomes, replicated in an independent cohort, and associated with a five-fold reduced risk of developing ALS. Variant IL18RAP 3′UTR reduces mRNA stability and the binding of RNA-binding proteins. Variant IL18RAP 3′UTR further dampens neurotoxicity of human iPSC-derived C9orf72-ALS microglia that depends on NF-κB signaling. Therefore, the variant IL18RAP 3′UTR provides survival advantage for motor neurons co-cultured with C9-ALS microglia. The study reveals direct genetic evidence and therapeutic targets for neuro-inflammation, and emphasizes the importance of non-coding genetic association studies.

**One Sentence Summary:** Non-coding genetic variants in IL-18 receptor 3’UTR decrease ALS risk by modifying IL-18-NF-κB signaling in microglia.

## Introduction

Genomic sequencing technologies facilitate the identification of variants in open reading frames (ORFs). Although allelic variants in non-coding regions are expected to be numerous ^1, 2^ they are largely overlooked.

Amyotrophic lateral sclerosis (ALS) is a fatal neurodegenerative syndrome, primarily affecting the human motor neuron system with a strong genetic predisposing component ^3, 4^. Thus far, mutations in approximately 25 protein-coding genes have been associated with ALS ^3, 5–7^. Hexanucleotide repeat expansion in an intronic sequence of the *C9orf72* gene is the most common genetic cause of ALS ^8–10^ and enrichment of variants was recently discovered in the CAV1 enhancer ^11^. However, non-coding nucleotide variants in ALS have yet to be systematically explored.

Burden analysis is a genetics approach that is based on the rationale that different rare variants in the same gene may have a cumulative contribution ^12^. Therefore, burden analysis allows the identification of genes containing an excess of rare and presumably functional variation in cases relative to controls. Although de novo mutations in non-coding regions were recently shown in family-based autism studies ^13^, variants in non-coding regions are not routinely included in rare-variant burden association studies. The application of burden analysis to non-coding regulatory variation is constrained by the availability of whole-genome sequencing (WGS) data, and the ability to recognize functional variants in non-coding regulatory regions, which is currently far less effective than for protein-coding genes.

MicroRNAs (miRNAs) are endogenous posttranscriptional repressors that silence mRNA expression through sequence complementarity. miRNA primarily act on 3′ untranslated regions (3′UTRs) ^14^, which are non-coding parts of messenger RNAs (mRNAs) and often regulate degradation and translation ^15^. miRNA dysregulation has been implicated in ALS pathogenesis, and ALS-associated RNA-binding proteins, TARDBP/TDP-43 and FUS, regulate miRNA biogenesis ^16–27^.

Microglia are the resident immune cells of the central nervous system and are the primary mediators of neuroinflammation in neurodegeneration ^28, 29^. In ALS, microglia induce motor neuron death via the classical nuclear factor-κappa B (NF-κB) pathway ^29–35^. One suggested mechanism for microglia-induced cytotoxicity is based on detection of extracellular TDP-43 aggregates and triggering of IL-1beta and Interleukin 18 (IL-18; also known as: IGIF/IL1F4/IL-1g) signaling ^31^. Accordingly, IL-18 levels are elevated in ALS patient tissues and biofluids ^36–39^, supporting a role for IL-18 signaling in the disease’s neuroinflammatory milieu ^40^. IL-18 also induces NF-κB signaling by binding and dimerising the two IL-18 receptor subunits, IL18RAP (also known as: AcPL/CD218b/IL-18R-Beta) and IL18R1 ^31, 40–48^. In turn, NF-κB contributes to microglial inflammation ^49, 50^, microglial-mediated motor neuron death ^51^ and to disease progression ^52, 53^.

Here, we identified rare variants in miRNAs and 3′UTR of mRNAs, and performed collapsed genetic analysis ^54^ to test if these regulatory RNAs are associated with ALS. We discovered an enrichment of rare variants in the IL18RAP 3′UTR and provide experimental evidence for their relevance to human ALS. Therefore, non-coding variant analysis reveals a genetic and mechanistic link for the IL-18 pathway in ALS and encourages systematic exploration of non-coding regions to uncover genetic mechanisms of disease.

## Results

To test whether genetic variations in non-coding regulatory regions are associated with ALS, we analyzed regions of interest in WGS data from the Project MinE ALS sequencing consortium ^55^ (Supplementary Fig. 1A,B and Supplementary Tables 1,2). The discovery cohort consisted of 3,955 ALS patients and 1,819 age- and sex-matched controls, for a total of 5,774 whole-genomes from the Netherlands, Belgium, Ireland, Spain, United Kingdom, United States, and Turkey (Project MinE Datafreeze 1). We performed a region-based burden test, in which rare genetic variants with minor allele frequencies (MAF) ≤0.01 are binned together, to weight their contribution to disease, in 295 non-coding 3′UTRs of candidate genes, linked to sporadic ALS via GWAS ^56^ or genes encoding RNA-binding proteins (Supplementary Table 3). In addition we tested all autosomal human-pre-miRNA genes (1,750 pre-miRNAs; miRBase v20 ^57^).

As a positive control, we also performed an association analysis of rare variants in the open reading frames of these 295 genes. For the proteins, we called variants that are predicted to cause frameshifting, alternative splicing, an abnormal stop codon, or a deleterious non-synonymous amino acid substitution that were detected in ≥ 3 of 7 independent dbNSFP prediction algorithms ^58^ (Fig. 1A and Supplementary Table 3). In total, 30,721 rare qualifying variants were identified (Supplementary Table 4). Optimized Sequence Kernel Association Test (SKAT-O) ^59^ identified a significant excess of deleterious minor alleles in the ALS genes *NEK1* (127 cases; 19 controls [3.21%; 1.04%]: P = 8×10^-7^; *P _corrected_*= 2.3×10^-4^), comparable with a reported prevalence of 3% ^60^, and in *SOD1* (36 cases [0.91%]; 0 controls: P = 2.6×10^-4^; *P _corrected_* = 3.73×10^-2^) ^61^, which is below the reported 2% prevalence ^5, 62^ (Fig. 1B, Supplementary Fig. 2A and Supplementary Table 5). Other known ALS genes did not reach statistical significance (Supplementary Table 3), consistent with reported statistical power limitations of Project MinE WGS data in assessing the burden of rare variants ^63^. Our analysis did not consider the *C9orf72* hexanucleotide (GGGGCC) repeat expansion region.

**Fig. 1.**
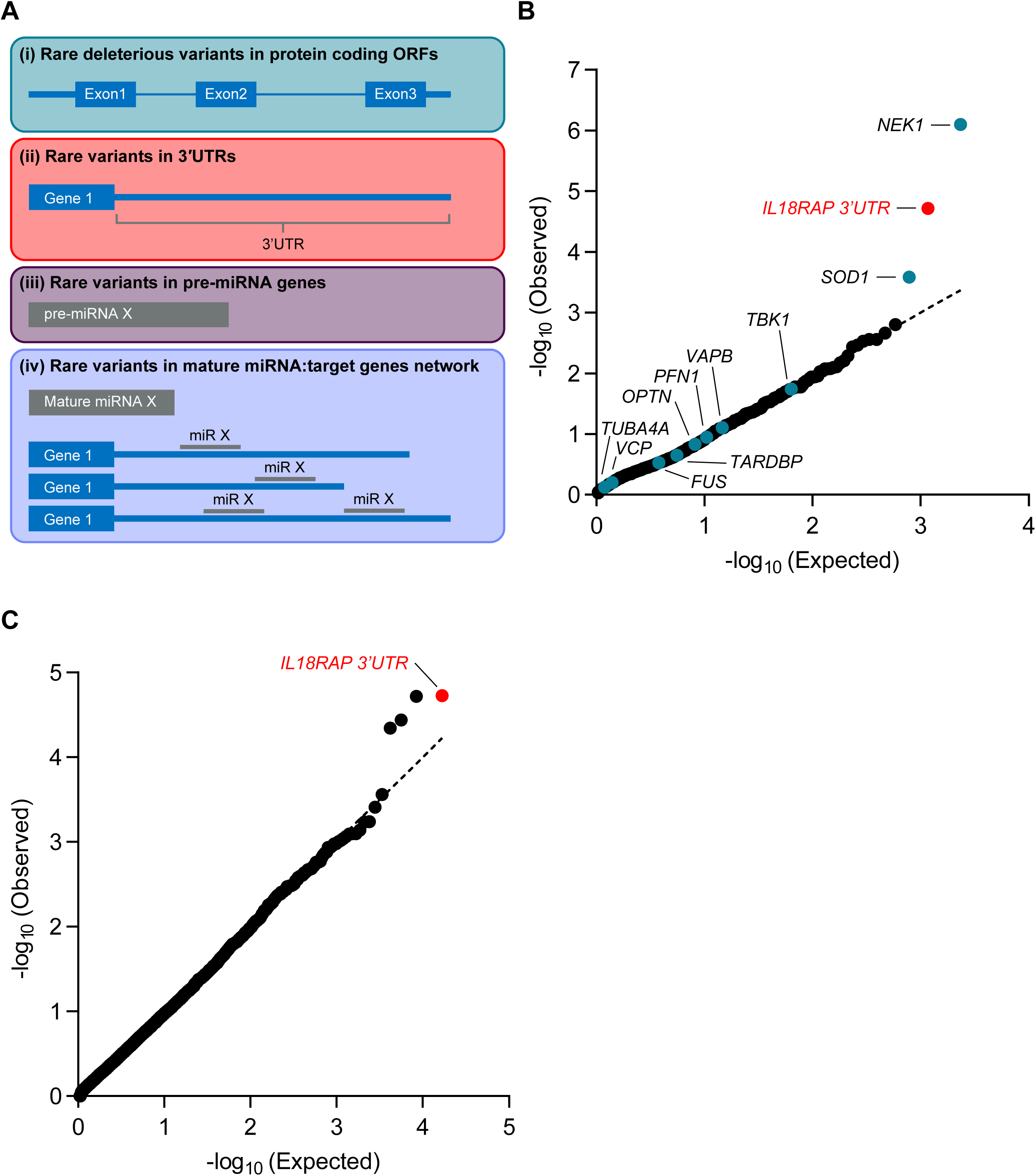
Region-based rare-variant association analysis reveals association of IL18RAP 3’UTR with ALS. **(A)** Diagram of study design. Collapsed region-based rare-variant (MAF ≤0.01) association analysis was performed on: (i) 295 candidate protein-coding genes (Supplementary Table 3), encoding for ALS-relevant proteins or proteins associated with miRNA biogenesis/activity. Variants were included if predicted to cause frameshifting, alternative splicing, an abnormal stop codon, or a deleterious non-synonymous amino acid substitution, in ≥ 3 of 7 independent dbNSFP prediction algorithms; (ii) variants in 3′-untranslated regions (3′UTRs) of the 295 genes (Supplementary Table 3); (iii) all known autosomal pre-miRNA genes in the human genome; and (iv) predicted networks, comprised of aggregated variants detected in a specific mature miRNA sequence and its cognate down-stream 3’UTR targets. **(B)** QQ plot of obtained and expected P-values for the burden of rare variants (log scale), gained by collapsed region-based association analysis of all genomic regions described in (A). Data were obtained from 3,955 ALS cases and 1,819 controls (Project MinE). Features positioned on the diagonal line represent results obtained under the null hypothesis. Open-reading frames of 10 known ALS genes (blue). IL18RAP 3′UTR (red). Genomic inflation λ = 1.2. **(C)** QQ plot of obtained and expected P-values for the burden of rare variants (log scale), gained by collapsed region-based association analysis for all known human 3’UTRs (RefSeq). The IL18RAP 3′UTR (red) is the most significant 3’UTR associated with ALS. P-values, calculated with Optimized Sequence Kernel Association Test, SKAT-O (genomic inflation λ = 0.97).

The burden of rare variants did not identify a disease association for any of the autosomal pre-miRNAs in the human genome, nor for any of the predicted genetic networks based on variants aggregated over specific mature miRNAs and their cognate downstream 3’UTR targets. This may be because the small size of miRNA genes makes genetic aggregation studies particularly challenging (Supplementary Fig. 2B,C).

When we tested the burden of variants in 3′UTRs, the strongest association found was for the 3′UTR of IL18RAP (Fig. 1B, Supplementary Fig. 2D and Supplementary Table 5). This association was higher than expected at random (P = 1.93×10^-5^, P *_corrected_* =5.41×10^-3^) and from the association gained for all protein-coding ALS genes in this cohort, with the exception of *NEK1*. Notably, the signal was more prevalent in controls [12/1819, 0.66%] relative to ALS patients [6/3955, 0.15%], indicating that these variants might act as protective variants against ALS.

IL18RAP 3′UTR also ranked as the top hit by three other algorithms – the Sequence Kernel Association Test (SKAT, P = 1.77×10^-5^; permutated P-value < 10^-4^), the Combined Multivariate and Collapsing (CMC, P = 8.78×10^-4^) or Variable Threshold (VT) with permutation analysis (permutated P-value = 1.75×10^-3^, suggesting that the association does not depend on a particular statistical genetics method (Supplementary Fig. 3A-C). Furthermore, when we tested the burden of variants in miRNA recognition elements (MREs) in 3’UTRs (variants that are potentially either abrogating conserved miRNA binding sites or creating new miRNA binding sites in 3’UTRs), the strongest association was also gained for the 3’UTR of IL18RAP (SKAT-O P-value = 3.42×10^-5^, Supplementary Fig. 3D, see Methods). A diagram of variants in IL18RAP 3’UTR is presented in Supplementary Fig. 3E and a description of IL18RAP 3′UTR variants in Supplementary Table 6. The top 10 principal components (PCs) of common variant-based ancestry information and sex were included as covariates in the SKAT-O, SKAT, CMC, and VT analyses.

In addition, genome-wide analysis of all known human 3’UTRs (16,809 3’UTRs from RefSeq ^64^) identified IL18RAP 3’UTR as the most significant 3’UTR associated with ALS in the Project MinE cohort (Fig. 1C).

Finally, we tested if different functional genetic classes were enriched overall for ALS risk/protection variants by testing the burden of rare variants in all genes pooled together. SKAT-O signal for open reading frames of 295 proteins, the 3’UTR of the same 295 genes, all autosomal pre-miRNA genes [miRBase v20; ^57^] or networks composed of all miRNA genes and their cognate set of downstream targets (TargetScan) were all not significant (P-values of 0.024, 0.59, 0.08, 0.58, respectively). Therefore, results from these burden tests do not implicate any of the functional class of genomic elements in ALS risk.

Because the number of ALS genomes was ∼2.17-fold larger than the number of controls, the data depict a 4.35- fold enrichment in the abundance of variants in controls over cases. IL18RAP 3′UTR potentially-protective variants reduced the disease odds ratio by five-fold (OR = 0.23; Fig. 2A), and was consistent across independent population strata (Fig. 2B), whereas *NEK1* and *SOD1* increased the disease odds ratio (OR = 3.14, 33.89, respectively; Fig. 2A).

**Fig 2.**
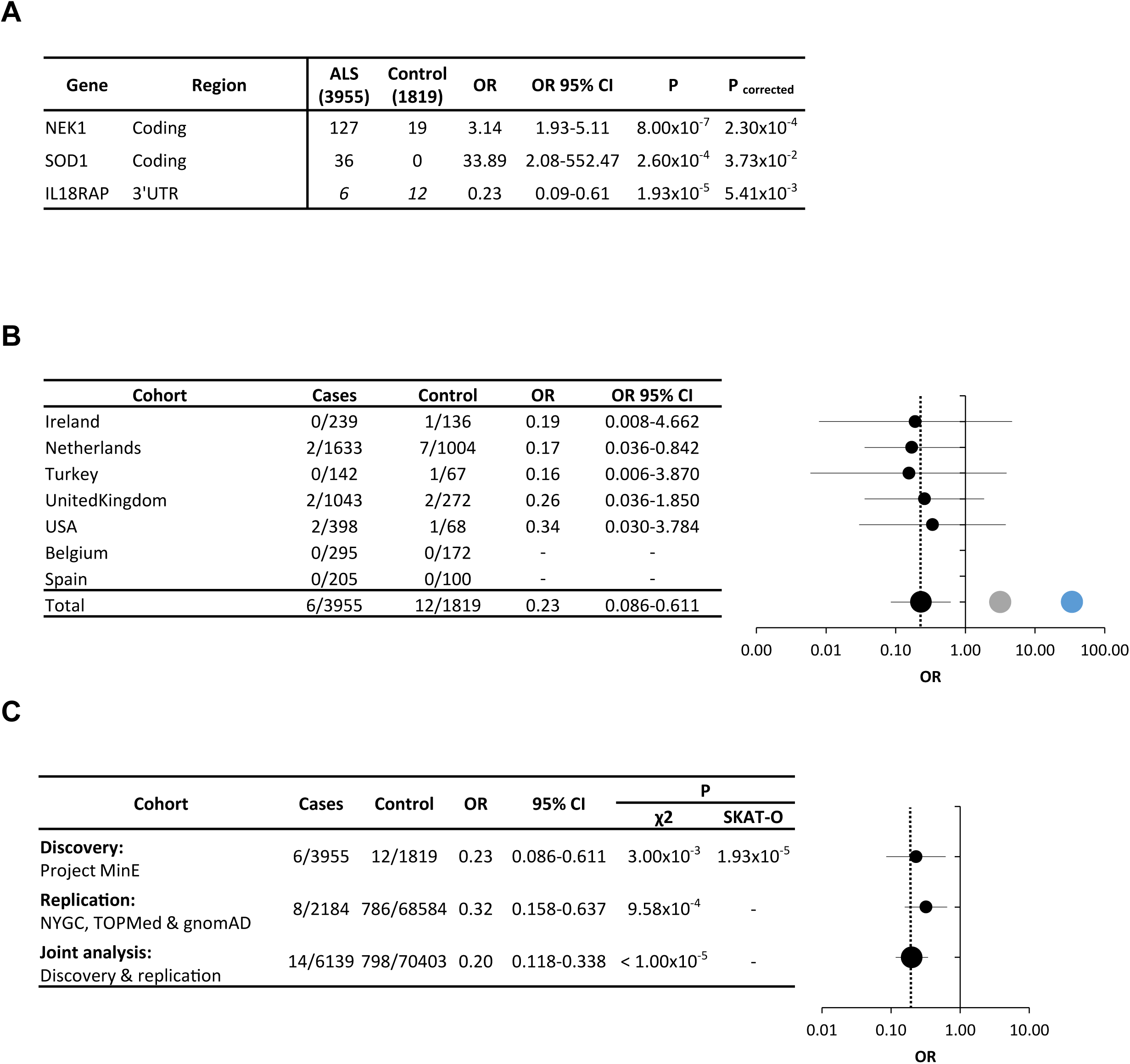
Odds of ALS are reduced with rare variants in the IL18RAP 3’UTR. **(A)** Odds ratio (OR) estimates with 95% confidence intervals (CI) for NEK1 (coding), SOD1 (coding) and IL18RAP (3’UTR). P-values corrected for false discovery rate (FDR). **(B)** Stratification of data pertaining to IL18RAP 3’UTR in seven geographically-based sporadic ALS sub-cohorts and forest plot (OR on log scale with whiskers for 95% CI). *NEK1* (grey) and *SOD1* (blue) signals are from combined data of all cohorts. Vertical dotted line denotes OR=0.23. **(C)** Stratification of IL18RAP 3’UTR variants data across discovery and replication cohorts and joint analysis thereof; Forest plot (OR on log scale with whiskers for 95% CI). Vertical dotted line denotes OR=0.2. P-values, calculated with SKAT-O or Chi-squared test with Yate’s correction.

To determine if the rare IL18RAP 3′UTR variants are depleted in another ALS cohort, we performed independent replication studies. Similar results for rare IL18RAP 3’UTR variants were reproduced in the New York Genome Center (NYGC) ALS Consortium cohort (2,184 ALS genomes), which was studied against: (i) 263 non-neurological controls from the NYGC; (ii) 62,784 non-ALS genomes from NHLBI’s Trans-Omics for Precision Medicine (TOPMed); and (iii) 5,537 non-ALS genomes from gnomAD. This replication effort yielded a joint analysis P-value = 9.58×10^-4^ (χ2 with Yate’s correction; OR=0.32; 95% CI: 0.16 – 0.64; Fig. 2C and Supplementary Table 7). Combining this cohort with our discovery cohort from Project MinE, yielded a superior joint P-value < 1.00×10^-5^ (χ2 with Yate’s correction; OR=0.20; 95% CI: 0.12 – 0.34; Fig. 2C). A meta-analysis of Project MinE datafreeze 1 and 2 ^7^, which consisted of 5,185 ALS patients and 2,262 age- and sex-matched controls, reproduced the initial signal (P-value = 7.6×10^-4^).

Together, IL18RAP 3’UTR sequence variants are associated with a lower risk of suffering from ALS, which is approximately one-fifth of the general population, although it did not reach conventional exome-wide multiplicity-adjusted significance threshold (α ≈2.6×10^-6^, ref. ^12^) in our study.

To investigate the source of the signal in the IL18RAP 3′UTR in a posthoc analysis, we divided the 11 variant nucleotides into two synthetic sets, of either nine singleton variants (9 variants / 3 controls / 6 patients) or two variants that were identified solely in controls (2 variants / 9 controls / 0 patients). While the signal of the nine singleton variants was not statistically significant, analysis of the two control variants, which were identified in multiple samples, derived an improved significance compared to the original signal (SKAT-O P-value = 4.36×10^-6^). Thus, these two rare variants (V1, Chr2:103068691 C>T; V3, Chr2:103068718 G>A) are likely central in generating the genetic association signal in IL18RAP 3′UTR.

Because of the enrichment of V1 and V3 at the proximal (5’) side of the IL18RAP 3’UTR, we tested if restricting burden analysis to the 5’ end of the 3’UTR, might boost the association signal. However, the P-values gained from the 3’UTRs proximal quadrant were comparable to that of the full 3’UTRs in the cohort of 295 3’UTRs (Wilcoxon matched-pairs P-value > 0.05, Cohen’s d effect size = 0.1, Supplementary Fig. 4A,B), suggesting that the apparent spatial distribution of variants in the case of IL18RAP 3’UTR is a particular case rather than part of a global pattern.

To determine the functional impact of the IL18RAP 3’UTR variants we analyzed IL18RAP expression in lymphoblastoid cell lines (LCLs) from the UK MNDA DNA Bank ^65^ that were derived from twelve different individuals: 4 healthy individuals (without ALS), carrying the canonical IL18RAP 3’UTR sequence (Control; Canonical IL18RAP 3’UTR); 4 sporadic ALS patients, carrying the canonical IL18RAP 3’UTR sequence (sALS; Canonical IL18RAP 3’UTR); two healthy individuals, carrying a variant form of IL18RAP 3’UTR (Control; VariantIL18RAP 3’UTR) and two sporadic ALS patients carrying a variant form of IL18RAP 3’UTR (sALS; VariantIL18RAP 3’UTR; see Supplementary Table 8 for list of variants).

ALS-derived LCLs carrying the canonical IL18RAP 3’UTR sequence expressed higher levels of IL18RAP (Fig. 3A,B). In addition, LCLs from both healthy and ALS individuals harboring the IL18RAP 3’UTR variant significantly down-regulated IL18RAP mRNA and protein expression (Fig. 3A,B and Data File S1). Phosphorylation of the nuclear factor kappa-light-chain-enhancer of activated B cells (p-NF-κB), an established intracellular effector downstream of IL-18 signaling, was similarly higher in the ALS LCLs with canonical IL18RAP 3’UTR and also significantly reduced in control and ALS LCLs harboring IL18RAP variants (Fig. 3C,D and Data File S1). Consistent results were obtained with C9orf72 hexanucleotide expansion ALS LCLs (Supplementary Fig. 5 and Data File S2). Accordingly, variants of IL18RAP 3’UTR reduced NF-κB activity, relative to the canonical 3’UTR in an NF-κB reporter assay in U2OS cells (Supplementary Fig. 6). Therefore, variant forms of IL18RAP 3’UTR correlate with reduced expression of the endogenous IL18RAP and reduced NF-κB signaling.

**Fig. 3.**
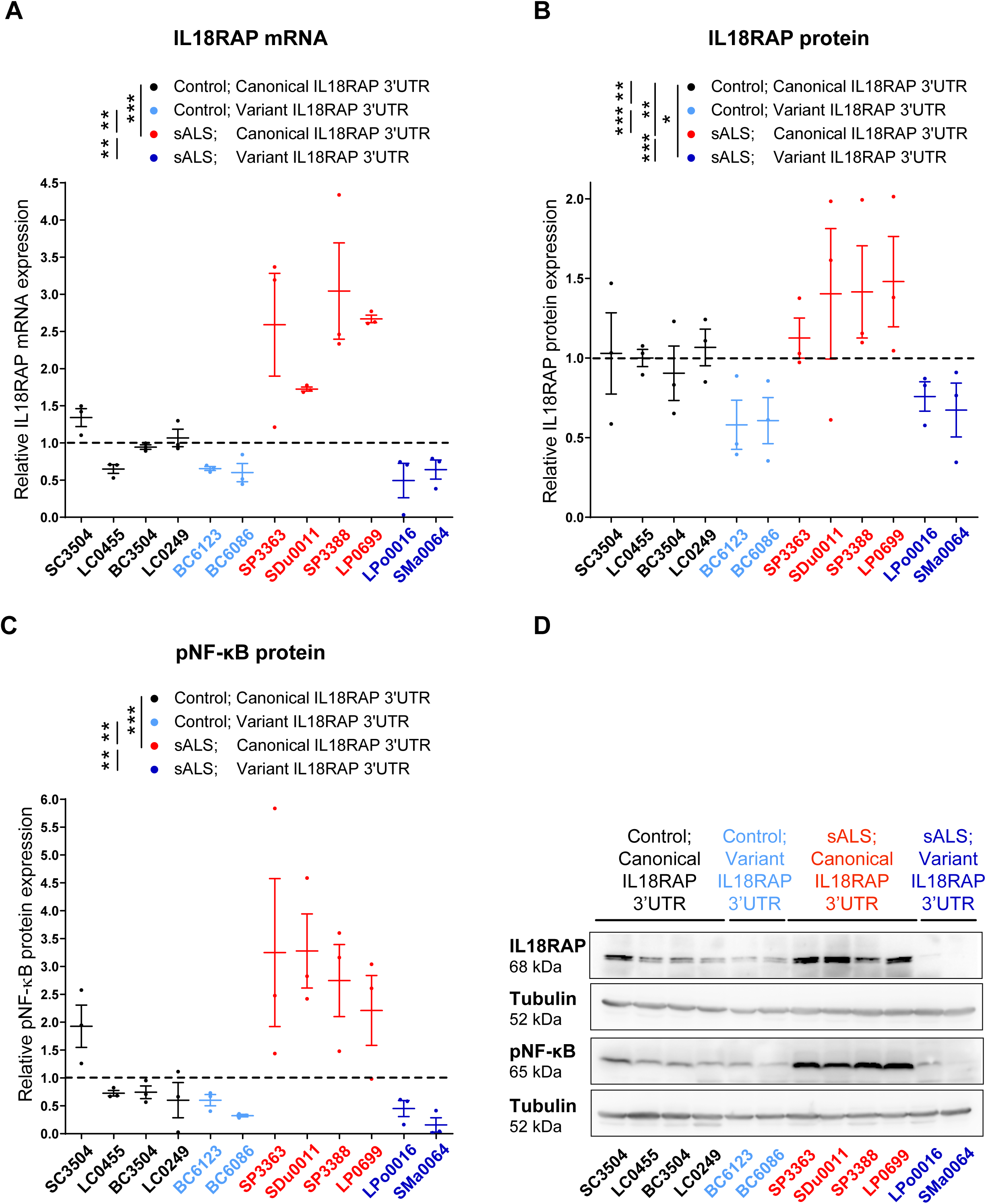
IL18RAP 3’UTR variant correlates with attenuated IL-18 - NF-κB signaling in human lymphoblastoid cells. **(A)** IL18RAP mRNA expression (qPCR normalized to IPO8 mRNA levels) and **(B)** IL18RAP or **(C)** p-NF-κB protein expression (Western blots, normalized to Tubulin). Scatter dot plot with mean and SEM. **(D)** Representative blots processed with anti-IL18RAP, anti p-NF-κB and anti-Tubulin antibodies. Extracts from twelve different human lymphoblastoid cell lines (listed in Supplementary Table 8): Four lines of healthy individuals (without ALS), carrying the canonical IL18RAP 3’UTR sequence (Control; Canonical IL18RAP 3’UTR, black); Four sporadic ALS patients, carrying the canonical IL18RAP 3’UTR sequence (sALS; Canonical IL18RAP 3’UTR, red); Two healthy individuals, carrying a variant form of IL18RAP 3’UTR (Control; Variant IL18RAP 3’UTR, light blue) and two sporadic ALS patients carrying a variant form of IL18RAP 3’UTR (sALS; Variant IL18RAP 3’UTR, navy blue). One-way ANOVA followed by Newman-Keuls multiple comparisons test, was conducted based on the mean value of three independent passages for each of the twelve human lymphoblastoid cell lines. * P<0.05; ** P<0.01; *** P<0.001.

To further establish the functional relevance of the IL18RAP 3’UTR variants, we edited the genome of human-induced pluripotent cells (iPSCs) donated by ALS patients with C9orf72 repeat expansion (^66^ NINDS/Coriell Code: ND10689, ND12099, see Supplementary Table 8) to include two point mutations that recapitulate the most prevalent variants (Chr2:103068691 C>T (V1) and Chr2:103068718 G>A (V3)) in the IL18RAP 3’UTR sequence (Fig. 4A). The resulting isogenic pair lines all carry C9orf72 repeat expansion and vary by only the presence of the canonical or a variant IL18RAP 3’UTR.

**Fig. 4.**
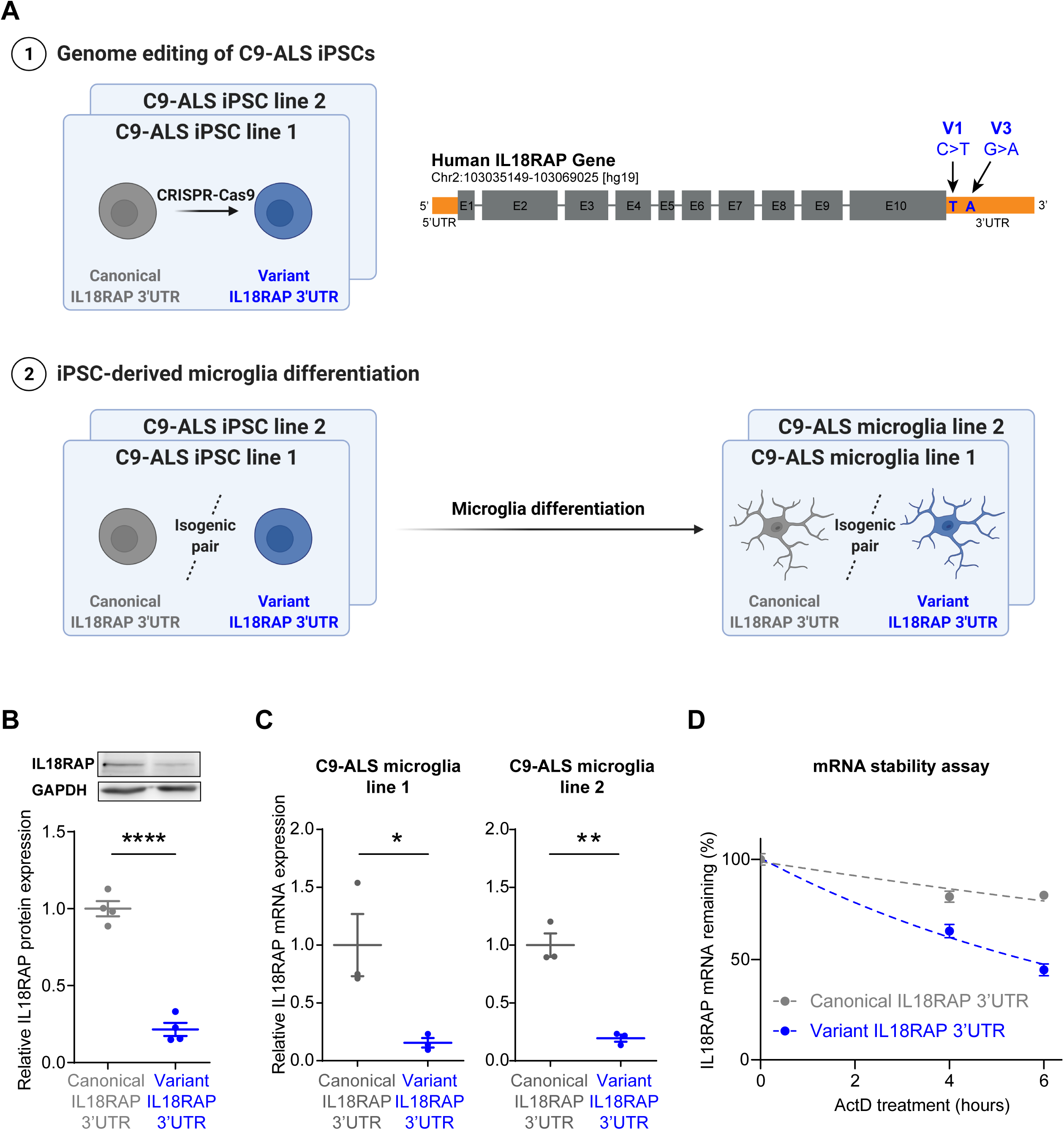
IL18RAP 3’UTR variant destabilizes IL18RAP mRNA in CRISPR-edited isogenic iPSC-derived microglia, with C9orf72 genetic background. (**A**) Diagram of experimental design. (1) Genome editing with CRISPR Cas9 of point mutations that recapitulate the most prevalent variants (Chr2:103068691 C>T (V1) and Chr2:103068718 G>A (V3)) in the IL18RAP 3’UTR sequence in human induced pluripotent stem cells (iPSCs) donated by ALS patients with a C9orf72 repeat expansion (^66^ NINDS/Coriell Code: ND10689, ND12099, Supplementary Table 8). The two independent isogenic pairs of cell lines both carry the C9orf72 repeat expansion and vary only by the presence of the canonical or a variant IL18RAP 3’UTR. (2) The four IL18RAP 3’UTR lines (two isogenic pairs) were differentiated into human microglia ^68^. Dot plots of IL18RAP (**B**) protein levels (by Western blot analysis, normalized to GAPDH, N=3, Data File S3) and (**C**) mRNA (by qPCR, normalized to IPO8 mRNA N=3) in differentiated human microglia. (**D**) IL18RAP mRNA degradation rate studied in human isogenic microglia at 0, 4 and 6 hrs after introduction of a transcriptional block with actinomycin D (7.5 μg/mL, Sigma-Aldrich A9415) (by qPCR, normalized to average of IPO8 and GAPDH mRNA expression, n = 4 independent wells per time point with two technical duplicates). Variant 3’UTR destabilizes the IL18RAP mRNA relative to the canonical sequence. Scatter dot plot with mean and SEM. Two sided t-test P-values * <0.05, ** <0.01, **** <0.0001.

We explored the receptive cell type involved in IL-18 receptor signaling by profiling dissociated mouse brain cells, namely, neurons, microglia, and astrocytes. Fluorescence cytometric gating on CD11b+ and CD45+ and IL18RAP (CD218b) revealed that IL18RAP is mainly expressed on microglia cells (Supplementary Fig. 7A-C). Although IL-18 and IL18RAP expression increases in ALS motor neurons (Supplementary Fig. 8A-C), our observations are consistent with the accepted notion that the role of IL-18 and other cytokines in disease heavily rests on a chronic inflammatory state established particularly by microglia ^67^.

Therefore, we next differentiated the isogenic IL18RAP 3’UTR lines into human microglia following the protocol of Haenseler et al. ^68^ (Fig. 4A). iPSC-derived microglia differentiation was validated by immunofluorescence staining of the microglial-specific marker, TMEM119 (Supplementary Fig. 9). In differentiated human microglia, we detected a ∼5-6 fold downregulation in the levels of the variant IL18RAP protein, as well as in the levels of the IL18RAP mRNA, relative to the canonical sequence of the isogenic line (Fig. 4B,C and Data File S3). Therefore, the variants at the 3’UTR regulate IL18RAP mRNA and protein expression and provide a conceivable explanation for the variant function in human C9-ALS microglia. Next, we investigated the molecular mechanism that controls the IL18RAP mRNA levels by performing an mRNA stability assay in human microglia. We measured an mRNA degradation rate that is twice as fast with the rare 3’UTR variants, relative to the canonical sequence, after inhibition of mRNA transcription by actinomycin D (Fig. 4D). Thus, the mechanism for reduced IL18RAP mRNA levels is associated with destabilization of IL18RAP mRNA via variants in the 3’UTR.

We sought the potential trans-acting factors that might differentially bind to the canonical and variant 3’UTRs. To this end, we performed RNA-pulldown assays and mass spectrometry on *in vitro* transcribed canonical and variant forms of the IL18RAP 3’UTRs, V1 and V3 (Fig. 5A diagram of exp. design). Mass spectrometry after pull-down identified 552 proteins with good confidence (passed all QC filters, found in 50% of the repeats in at least one experimental group, and were represented by at least 2 unique peptides, Supplementary Table 9), that were enriched in comparison to the negative control. Principal component analysis demonstrated a clear separation of proteomes bound by the canonical and variant IL18RAP 3’UTRs (Fig. 5B and Supplementary Table 9). Gene set enrichment analysis (GSEA) revealed a reduction in the association of double-stranded RNA (dsRNA) binding proteins, to V1 IL18RAP 3’UTR, relative to the canonical 3’UTR (ELAVL1/Hur; PRKRA, EIF2AK2/PKR; ADAR; ADARB1; ILF2; ILF3; DHX9; DHX58; DDX58, Fig. 5C,D,E and Supplementary Table 10). These dsRNA binding proteins were reported in other contexts to play roles in controlling the stability of mRNA ^69–75^, consistent with the observed changes to IL18RAP mRNA stability. A similar analysis of the V3 variant was unproductive (Supplementary Fig. 10A).

**Fig. 5.**
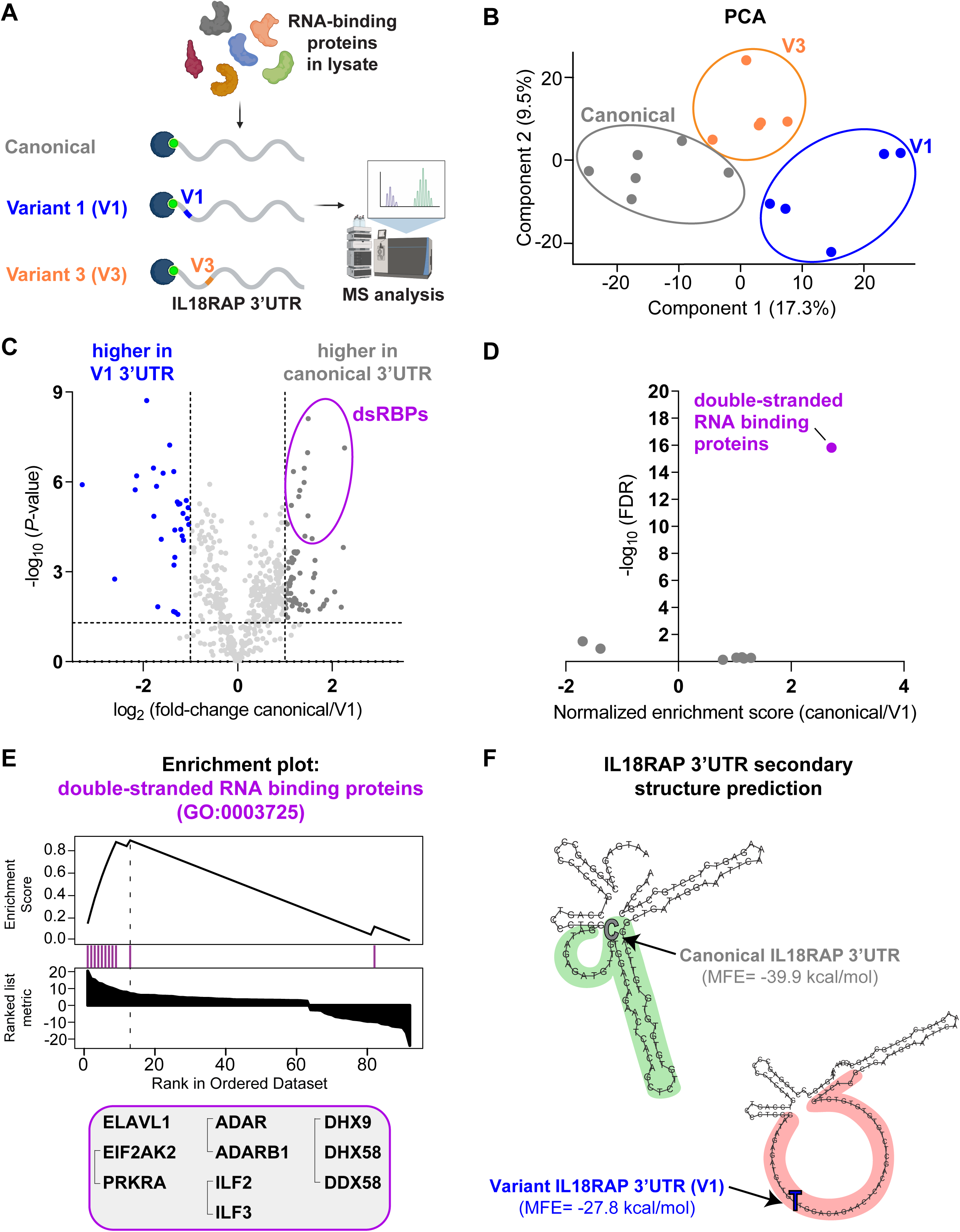
Reduced association of double-stranded RNA binding proteins to variant IL18RAP 3’UTR. **(A)** Diagram of mass spectrometry of RNA binding proteins pulled-down by IL18RAP 3’UTR sequences (canonical, V1 and V3). **(B)** Principal-component analysis (PCA) of IL18RAP 3’UTR-associated proteomes pulled down by the canonical (grey, N=6 experimental repeats), V3 (orange, N=5), and V1 (blue, N=5) biotin-tagged, in-vitro transcribed oligos. **(C)** Volcano plot of protein abundance associated with the canonical relative to variant (V1) IL18RAP 3’UTR (x-axis log2 scale), analyzed by MS. Y-axis depicts P-values (−log10 scale). Proteins significantly enriched in association with canonical/variant 3’UTR are colored (grey/blue). Double-stranded RNA-binding proteins (dsRBPs) are demarcated by a purple oval. Features above the horizontal dashed line demarcate proteins with adjusted p < 0.05, in student’s t-test with FDR correction to multiple hypotheses. Vertical dashed lines are of 2 or ½ fold change. A non-significant data point of KIF13B (P-value = 0.08) is not shown for clarity of the illustration (Supplementary Table 9) **(D)** Volcano plot of normalized enrichment score of the Gene Ontology (GO) molecular function gene sets from GSEA analysis of differentially expressed proteins (canonical vs. V1 IL18RAP 3’UTR). Reduced association of double-stranded RNA binding proteins (GO:0003725; purple) with V1 IL18RAP 3’UTR, relative to the canonical 3’UTR. All gene sets are described in Supplementary Table 10. **(E)** Profile of GSEA enrichment score and positions of the 10 double-stranded RNA binding proteins (purple) within all differentially expressed proteins, ranked from most enriched in canonical 3’UTR to most depleted protein (Supplementary Table 10; WebGestalt ^104^). **(F)** Prediction of 3’UTR secondary structure by RNA Fold ^105^ suggests a more stable dsRNA structure of canonical 3’UTR (green), with lower minimum free energy (MFE) than that of the sequence harboring a V1 variant (red).

In accordance, RNA Fold analysis predicted that the canonical 3’UTR sequence consists of a more stable dsRNA structure than the V1 variant sequence (minimum free energy (MFE) of canonical and variant IL18RAP 3’UTR, - 39.9 kcal/mol and -27.8 kcal/mol, respectively) (Fig. 5F and Supplementary Fig. 10B). In light of these results, we propose that variant-dependent changes to the secondary structure of IL18RAP 3’UTR attenuate the binding of one or more of the dsRNA proteins and may be involved in controlling the stability of IL18RAP mRNA.

To study the potential protective impact of IL18RAP 3’UTR mutations, we performed survival analyses in a coculture system of human iPSC-derived isogenic IL18RAP 3’UTR microglia (on a *C9orf72 repeat expansion background)* with human iPSC-derived lower motor neurons (i^3^LMNs; healthy, non-ALS, ^76^). Time-lapse microscopy was used to quantify motor neuron survival after microglia activation with a cocktail of LPS and the cytokine IL-18 (experimental design, Fig. 6A). Motor neuron survival was significantly improved in the presence of microglia harboring the IL18RAP 3’UTR variants relative to microglia harboring the canonical IL18RAP 3’UTR (two independent isogenic pairs, based on independent patient C9orf72 lines, n=3 independent differentiation procedures from different passages per line, with 3-8 co-culture wells per passage; Fig. 6B-D, Supplementary movie and Data File S4). Based on these studies, we conclude that rare variants of IL18RAP 3’UTR increase *C9orf72* microglia-dependent motor neuron survival and hence convey a protective property.

**Fig. 6.**
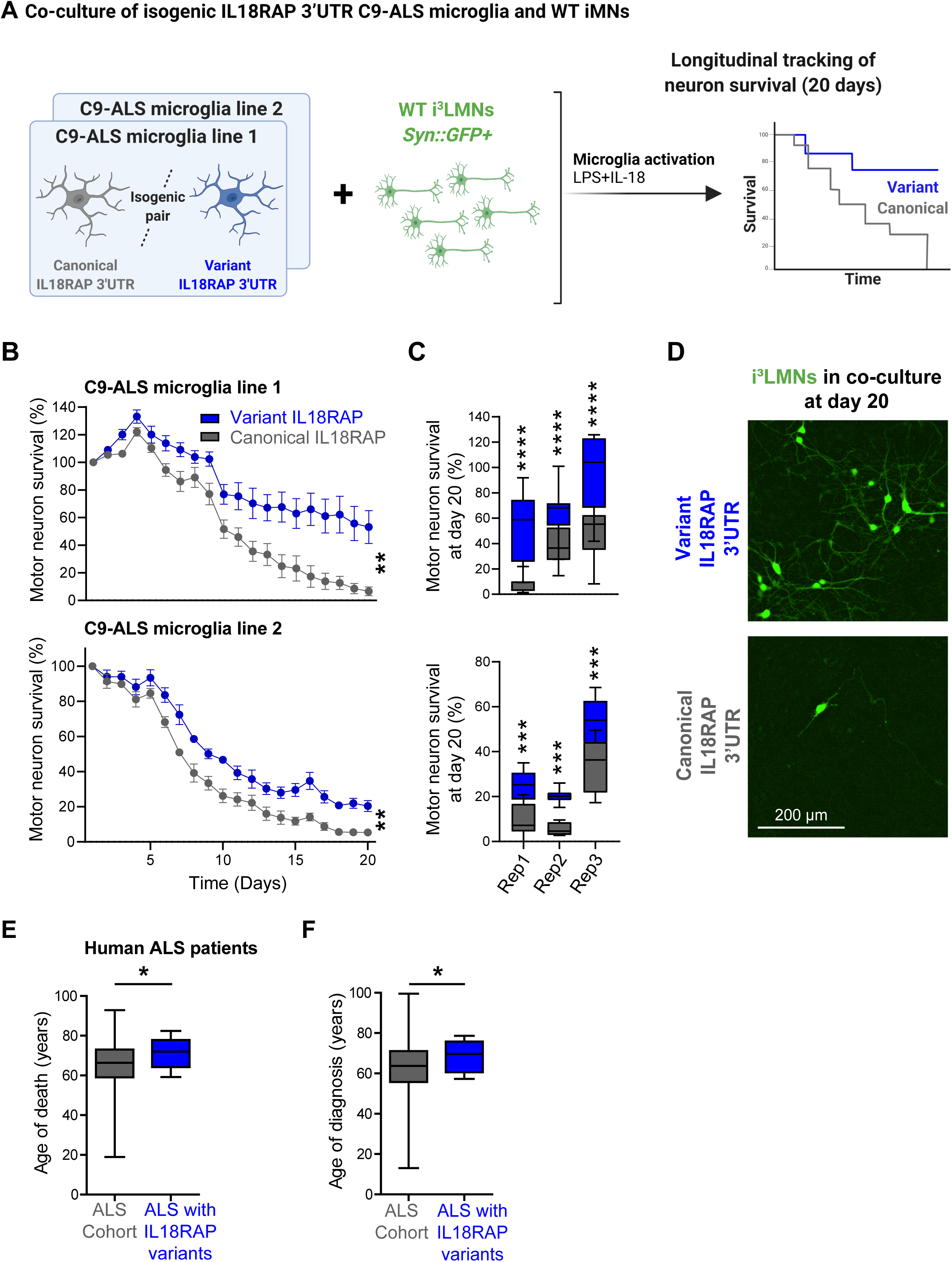
Variant IL18RAP 3’UTR are protective in human microglia and in patients with ALS. **(A)** Diagram of experimental design. Co-culture of human iPSC-derived transcription factor-induced motor neurons (i^3^LMNs) that express GFP driven by the synapsin (Syn) promoter (healthy, non-ALS, ^76^) and human iPSC-derived isogenic IL18RAP 3’UTR microglia (on a *C9orf72 repeat expansion background)*. Time-lapse microscopic analyses of i^3^LMNs survival, after microglia activation with a cocktail of LPS and the cytokine IL-18. **(B,C)** i^3^LMNs survival over 20 days in the presence of microglia harboring variant (blue) or canonical (grey) IL18RAP 3’UTR (two independent isogenic pairs, based on independent patient C9orf72 lines, n=3 independent differentiation procedures from different passages per line, with 3-8 co-culture wells per passage). **(B)** Survival plot of i^3^LMNs in a representative experiment for each isogenic pair (Two-way ANOVA) and **(C)** Box plot depicting the percentage of i^3^LMNs survival on day 20 of co-culture, median, upper and lower quartiles of all experiments. Two independent isogenic pairs, based on independent patient C9orf72 lines, n=3 independent differentiation procedures from different passages per line, with 3-8 co-culture wells per passage. Two-way ANOVA followed by Tukey’s multiple comparison test. **(D)** Representative micrographs of fluorescent i^3^LMNs after 20 days of culture with C9-ALS microglia. **(E)**. Association of age of death (9 patients with protective 3’UTR variants /4263 patients with available phenotypic data in Project MinE and NYGC cohorts, or **(F)** age of diagnosis (8/4216 patients). IL18RAP variant is associated with delayed age of death (+6.1 years, Permutation P-value = 0.02, Cohen’s d effect size = 0.65) and age of diagnosis (+6.2 years, Permutation P-value = 0.05, Cohen’s d effect size = 0.62), relative to the mean age of all Project MinE and NYGC ALS patients. Box plots depicting median, upper and lower quartiles, and extreme points. * P<0.05, ** P<0.01, *** P<0.001, **** P<0.0001.

To determine whether the mutant IL18RAP 3’UTR is also protective in human patients with ALS, we tested the association between age of diagnosis and age of death in ALS patients harboring canonical or variants of the IL18RAP 3’UTR. Of 4216 patients for whom data on the age of diagnosis was available (Project MinE and NYGC cohorts), 8 harbored IL18RAP 3’UTR variants. Of 4263 patients for whom the age of death was available, 9 harbored IL18RAP 3’UTR variants. IL18RAP 3’UTR variants are expected to be depleted in ALS genomes, nonetheless, in those extremely rare patients harboring IL18RAP 3’UTR variants, these were associated with an older age of death and an older age of diagnosis. On average, the age of death was higher by 6.1 years after the average for patients with canonical Il18RAP 3’UTR (Permutation P-value = 0.02, Cohen’s d effect size = 0.65; Fig. 6E and Supplementary Table 11), and the age of diagnosis was higher by 6.2 years after the average for patients with canonical IL18RAP 3’UTR (Permutation P-value = 0.05, Cohen’s d effect size = 0.62; Fig.6F and Supplementary Table 11). Thus, variants in IL18RAP 3’UTR are protective against ALS in a tissue culture model and correlate with survival advantage for patients suffering from the disease.

To study the role of NF-κB signaling in our system, we analyzed NF-κB phosphorylation and the impact on the transcriptome after microglia activation (Fig. 7A). Western blot analysis revealed reduced levels of phospho-NF-κB in variant IL18RAP 3’UTR relative to isogenic control (Fig. 7B and Data File S5). Reduced phosphorylation is associated with decreased nuclear localization and transcriptional activity of NF-κB ^77–80^. In parallel, we conducted a next-generation sequencing study (Supplementary Table 12, Gene Expression Omnibus accession number: GSE186757) of the differentially expressed transcriptomes in microglia harboring variant vs. canonical IL18RAP 3’UTR. Over-representation analysis (ORA) of differentially expressed genes (DEGs) revealed downregulation of the NF-κB signaling pathway in microglia harboring the variant IL18RAP 3’UTR (KEGG Pathway enrichment results: Ratio = 3.77, FDR P-value = 7.34×10^-6^; Gene Ontology Biological Process enrichment results: Ratio = 3.48, FDR P-value = 3.70×10^-12^, Fig. 7C,D and Supplementary Table 13). In addition, an unsupervised study of NF-κB pathway mRNAs (GO:0007249) demonstrated broad downregulation of pathway-associated mRNAs in microglia with the variant IL18RAP 3’UTR, relative to the isogenic control (Fig. 7E). Therefore, microglia’s NF-κB transcriptomic signature depends on signaling via the IL-18 receptor and is attenuated by protective IL18RAP 3’UTR variants.

**Fig. 7.**
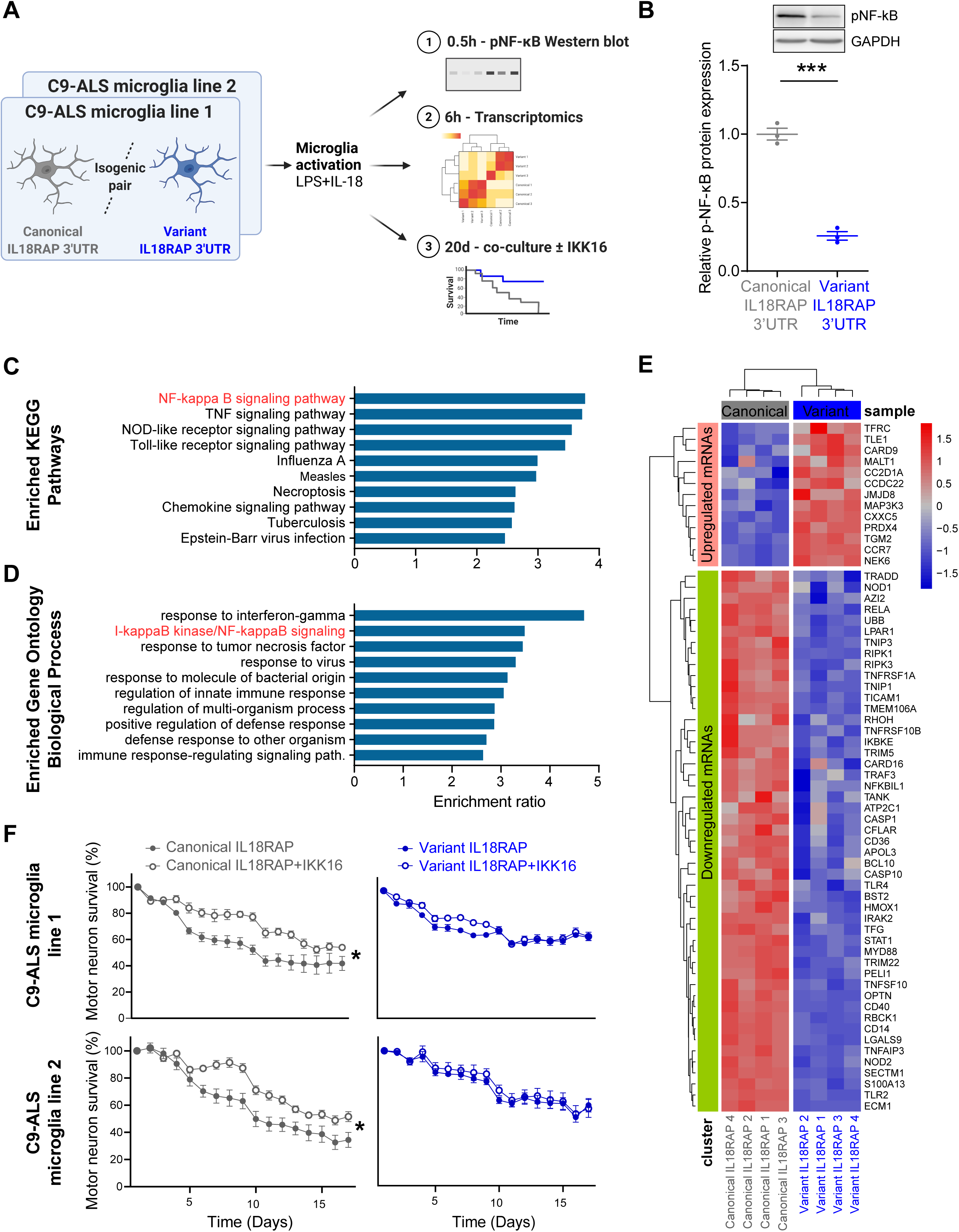
Variant IL18RAP 3’UTR dampens neurotoxic NF-κB signaling in human microglia. **(A)** Diagram of experimental design. Four IL18RAP 3’UTR lines (two isogenic pairs) were differentiated into human microglia ^68^ and analyzed for phosphorylated NF-κB protein levels, transcriptomics, and neuronal survival in co-culture with/without IKK16, following activation with a cocktail of LPS and the cytokine IL-18, for 0.5h, 6h and 20 days, respectively. **(B)** Western blot analysis revealed reduced levels of phosphorylated NF-κB in variant IL18RAP 3’UTR relative to isogenic control. Scatter dot plot with mean and SEM (Two sided t-test P-value *** <0.001, N=3, Data File S5). mRNA extracted from human microglia was subjected to a next generation sequencing study with downstream bioinformatics studies **(C-E)**. Over-representation analysis (ORA) within **(C)** KEGG Pathways and **(D)** Gene Ontology biological processes, of the differentially expressed transcriptome in microglia harboring variant vs. canonical IL18RAP 3’UTR. Bar graph depicting the Ratio of enrichment for significantly enriched pathways (FDR ≤ 0.05) are shown (Supplementary Table 13; WebGestalt ^104^). **(E)** Unsupervised study of the NF-κB transcriptomic signature (GO:0007249 pathway-associated mRNAs) in microglia with the variant relative to the isogenic canonical IL18RAP 3’UTR. **(F)** Time-lapse microscopic analyses of co-cultured human i^3^LMNs (healthy, non-ALS, ^76^) with human iPSC-derived isogenic IL18RAP 3’UTR microglia (on a *C9orf72 repeat expansion background)*, activated with a cocktail of LPS and the cytokine IL-18, without (carrier alone, DMSO), or with IKK16 (200nM), a selective IκB kinase (IKK) inhibitor that inhibits NF-κB signaling ^81^. IKK16 significantly ameliorates motor neuron death, relative to control only in the context of canonical IL18RAP 3’UTR, but did not further contribute to rescue in human microglia with the protective variant IL18RAP 3’UTR (two independent isogenic pairs, based on independent patient C9orf72 lines with 3-8 co-culture wells per line). Two-way ANOVA P-value * <0.05.

To test a plausible neurotoxic role for NF-κB downstream of the IL-18 receptor in this system, we next performed a co-culture survival assay with or without IKK16, a selective IκB kinase (IKK) inhibitor that inhibits NF-κB signaling ^81^. In human microglia with the canonical IL18RAP 3’UTR, IKK16 significantly ameliorated motor neuron toxicity, relative to control (carrier alone, Fig. 7F). However, in human microglia with the protective variant IL18RAP 3’UTR, inhibition of NF-κB had no effect (two independent isogenic pairs, based on independent patient C9orf72 lines with 3-8 co-culture wells per line, Fig. 7F). This suggests that NF-κB neurotoxic function resides epistatically downstream of IL18RAP in human microglia. Together, rare variants in IL18RAP 3’UTR diminish NF-κB signaling, thus increasing *C9orf72* microglia-dependent motor neuron survival.

## Discussion

Data from the Project MinE and NYGC ALS consortia provide unprecedented opportunities for investigating the role of the non-coding genome in ALS. By performing rare variant aggregation analysis in regulatory non-coding regions, we demonstrate that variants in the 3′UTR of IL18RAP are enriched in non-ALS genomes, indicating that these are relatively depleted in ALS. IL18RAP 3′UTR variants reduced the chance of developing ALS five-fold, and delayed onset and therefore age of death in people with ALS.

These protective variants recall other protective variants that have been reported in the past in protein-coding regions in Alzheimer’s disease ^82–85^ and implicated in ALS as well ^86, 87^. In addition, deleterious variants were suggested in VEGF promoter/5’UTR and within CAV1/CAV2 enhancers ^11, 88^. However, the 3′UTR of IL18RAP is a protective non-coding allele associated with a neurodegenerative disease.

Elevated levels of the cytokine IL-18 were reported in tissues and biofluids of ALS patients ^36–38^. Accordingly, we reveal the upregulation of endogenous IL18RAP in sporadic and C9orf72 lymphoblastoid cells. In addition, we demonstrate the downregulation of IL18RAP expression in lymphoblastoid cells harboring variant versions of the IL18RAP 3’UTR.

We elucidated the regulatory changes affected by the IL18RAP 3′UTR variants by showing destabilization of the IL18RAP mRNA and downregulation of IL18RAP mRNA levels. Sequence analysis suggests that at least one variant (V1) potentially reduced the propensity of the 3’UTR to form a double-stranded secondary structure. Accordingly, unbiased proteomics demonstrated that the 3’UTR harboring the variant fails to bind dsRNA binding proteins that are known to stabilize mRNAs. Together, this supports a mechanism for reduced IL18RAP signaling involving changes to mRNA stability and differential binding of stabilizing RNA-binding proteins.

Neuro-inflammation is prevalent in neurodegeneration, including in ALS ^89^, and is often characterized by the activation of microglia ^29–35^. The cytokine, IL-18, is part of this neuro-inflammatory milieu, promoting receptor subunit (IL18RAP, IL18R1) dimerization on the membrane of cells ^40^, and activating intracellular signaling cascades, including NF-κB.

By CRISPR editing of two independent C9orf72 lines, from one female and one male patient, we created isogonic IL18RAP 3’UTR cell lines (canonical or harboring V1 and V3 variants), at the endogenous gene locus. By differentiating these lines to human microglia, we demonstrated that variants downregulated IL18RAP mRNA and protein expression. In addition, tracking of human (wild-type) motor neuron survival, in co-culture with microglia, over 21 days, demonstrated the neuroprotective effect of microglia carrying the variant IL18RAP 3’UTR.

Finally, we demonstrate that variant IL18RAP 3’UTR attenuates NF-κB signaling in lymphoblastoid cells and in microglia. Unbiased next-generation sequencing of microglia RNA demonstrated broad transcriptomic changes, typical of reduced NF-κB signaling. In addition, inhibition of NF-κB was able to ameliorate motor neuron death when co-cultured with microglia harboring the canonical IL18RAP 3’UTR. However, inhibition of NF-κB was not further protective if microglia with variant IL18RAP 3’UTR were present, suggesting an epistatic relationship, whereby IL18RAP is upstream of NF-κB in this system. We conclude that IL18RAP acts in microglia and controls the cytotoxicity conveyed to motor neurons, at least in human C9orf72 types of disease.

The discovery of functional, disease-modifying IL18RAP 3’UTR variants underscores the need to explore the role of additional non-coding genomic regions in ALS. One limitation of our study is that IL18RAP 3’UTR signal did not reach the conventional exome-wide multiplicity-adjusted significance threshold (α ≈2.6×10^-6^, ref. ^12^). However, IL18RAP 3’UTR signal is comparable to that of protein-coding ALS-causing genes, such as *SOD1* and *NEK1.* Furthermore, the key findings were reproduced in a genome-wide study of all human 3’UTRs and in an independent replication study. Limitations in the statistical power might have prevented the discovery of other non-coding variants and may be overcome with larger ALS and control cohorts, which are not currently available. Additionally, we have focused our tissue culture studies on human C9orf72 microglia. Therefore, the involvement of IL18RAP 3’UTR in other ALS-associated genetic backgrounds remains to be experimentally explored, as is the relevance to other neurodegenerative diseases. Finally, the mechanism underlying IL18RAP dose sensitivity is not fully understood. While we provide evidence that variant IL18RAP 3’UTR endows neuroprotection via dampening of microglia-dependent neurotoxicity, additional studies should explore the degree to which other cell types, such as motor neurons and astroglia, are involved.

In summary, we have identified the IL18RAP 3′UTR as a non-coding genetic disease modifier by burden analysis of WGS data using ALS case-control cohorts. We show that IL-18 signaling modifies ALS susceptibility and progression, delineating a neuro-protective pathway and identifying potential therapeutic targets for ALS. Whereas the 3′UTR of IL18RAP is a protective non-coding allele associated with a neurodegenerative disease, the increasing wealth of WGS data in Project MinE, NYGC and elsewhere, indicates that the exploration of non-coding regulatory genomic regions should reveal further disease-relevant genetic mechanisms.

## Methods

### Human genetic cohorts

All participants contributed DNA after signing informed consent at the submitting sites. Human materials were studied under approval of the Weizmann Institute of Science Institutional Review Board (Weizmann IRB: 1039-1).

Discovery cohort: Project MinE ALS sequencing consortium Datafreeze 1 includes 3,955 ALS patients and 1,819 age- and sex-matched controls, free of any neurodegenerative disease, for a total of 5,774 quality control (QC) passing whole-genomes, from the Netherlands, Belgium, Ireland, Spain, United Kingdom, United States, and Turkey. Rare variant association in cases versus controls was evaluated for regions of interest, when we could identify ≥2 variants per region, by SKAT-O, SKAT, CMC, and VT in RVTESTS environment ^90^, with sex and the top 10 principal components (PCs) as covariates. To construct the PCs of the population structure, an independent set of ∼450,000 SNPs was sampled from WGS, (MAF ≥0.5%) followed by LD-pruning. Rare genetic variants were included based on minor allele frequencies (MAF) ≤0.01 within the controls in the current data set.

Replication cohorts: Utilized for testing rare variant alleles (MAF < 0.01) in human IL18RAP 3’UTR (GRCh37/hg19 chr2:103068641-103069025 or GRCh38 chr2:102452181-102452565) from Project MinE datafreeze 2: ∼1300 European heritage ALS genomes without middle eastern (Turkish and Israelis) genomes. The New York Genome Center (NYGC) ALS Consortium (2,184 ALS Spectrum MND and 263 non-neurological control genomes from European/Americas ancestries), NHLBI’s Trans-Omics for Precision Medicine (TOPMed; 62,784 non-ALS genomes) and gnomAD (5,537 non-ALS genomes; Europeans, non-Finnish, non-TOPMed). Joint analysis in replication cohort was performed by Chi-square test with Yate’s correction. Meta-analysis was not possible because TOPMed and gnomAD covariate information is not available.

### Quality control procedures in Project MinE genomics

Sample selection, data merging, and sample- and variant level quality control procedures for Project MinE ALS sequencing consortium genomes are described in full previously ^63^. Briefly, 6,579 Project MinE ALS sequencing consortium whole genomes were sequenced on Illumina HiSeq2000 or HiSeqX platforms. Reads were aligned to human genome build hg19 and sequence variants called with Isaac Genome Alignment Software and variant caller ^91^. Individual genomic variant call format files (GVCFs) were merged with ‘agg’ tool: a utility for aggregating Illumina-style GVCFs. Following completion of the raw data merge, multiple QC filtering steps were performed: (i) setting genotypes with GQ<10 to missing; (ii) removing low-quality sites (QUAL< 30 and QUAL< 20 for SNPs and indels, respectively); (iii) removing sites with missingness > 10%. (iv) Samples excluded if deviated from mean by more than 6SD for total numbers of SNPs, singletons and indels, Ti/Tv ratio, het/hom-non-ref ratio, and inbreeding (by cohort). (v) missingness > 5%, (vi) genotyping-sequence concordance (made possible by genotyping data generated on the Illumina Omni 2.5M SNP array for all samples; 96% concordance), (vii) depth of coverage, (viii) a gender check (to identify mismatches), (ix) relatedness (drop samples with >100 relatedness pairs). (x) Related individuals were further excluded until no pair of samples had a kinship coefficient > 0.05. (xi) missing phenotype information. Following QC, 312 samples with expended/inconsistent *C9orf72* status were omitted from further analysis. A total of 5,774 samples (3,955 ALS patients and 1,819 healthy controls) passed all QC and were included in downstream analysis. Per-nucleotide site QC was performed on QC-passing samples only, for Biallelic sites: variants were excluded from analysis based on depth (total DP < 10,000 or > 226,000), missingness > 5%, passing rate in the whole dataset < 70%, sites out of Hardy–Weinberg equilibrium (HWE; by cohort, controls only, p < 1×10^-6^) and sites with extreme differential missingness between cases and control samples (Overall and by cohort, p < 1×10^-6^). Non-autosomal chromosomes and multiallelic variants were excluded from analysis.

### Selection of regions of interest

Discontinuous regions of interest approximating in total ∼5Mb, include coding sequences and 3′ untranslated regions (3′UTRs) of 295 genes (Supplementary Table 3) encoding for proteins that were: (i) previously reported to be associated with ALS, (ii) RNA-binding proteins including miRNA biogenesis or activity factors [UCSC gene annotation; ^92^]. In addition to (iii) all 1,750 autosomal human pre-miRNA genes [miRBase v20; ^57^]. In addition, genome-wide analysis of all known human 3’UTRs (RefSeq ^64^). Variants in regions of interest were extracted from Project MinE ALS sequencing consortium genomes using vcftools ^93^ according to BED file containing genomic coordinates of interest (hg19) ±300 bp that ensures covering splice junctions and sequence (Supplementary Table 14).

### Annotation and burden analysis

After quality control and extraction of regions of interest, we performed functional annotation of all variants. Indels were left-aligned and normalized using bcftools and multiallelic sites were removed. For variant annotation we developed a pipeline that calculates the impact of genetic variation in coding regions as well as in 3’UTR and miRNA regions, using ANNOVAR ^94^. The frequency of the variants in the general population was assessed by screening the 1000 Genomes Project, the Exome Aggregation Consortium (ExAC), and NHLBI Exome Sequencing Project (ESP). For protein-coding ORFs, association analysis of deleterious rare variants was performed, i.e., frameshift variants, deviation from canonical splice variant, stop gain/loss variants, or a non-synonymous substitution, as predicted by at least three prediction programs (SIFT, Polyphen2 HVAR, LRT, MutationTaster, MutationAssessor, FATHMM, MetaLR) in dbNSFP environment [v2.0; ^58^].

Non-coding sequence burden analysis included (i) 3′UTRs, (ii) variants in miRNA recognition elements (MREs) in 3’UTRs (Supplementary Table 3): Variants that impaired conserved-miRNA binding sites in 3’UTRs (predicted loss of function) were called by TargetScan [v7.0; ^95^]. Newly created miRNA binding sites in 3’UTRs (predicted gain of function) were called by textual comparison of all possible mutated seeds around a variant to all known miRNA seed sequences in the genome, (iii) all human pre-miRNAs (mirBase v20 ^57^) and (iv) miRNAs:target gene networks: mature miRNA sequences (mirBase v20 ^57^) and cognate targets within the 3’UTRs (Supplementary Table 3). Variant annotation scripts are available at GitHub: https://github.com/TsviyaOlender/Non-coding-Variants-in-ALS-genes-

### Mammalian Cell Cultures

Lymphoblastoid cell lines (LCLs) from the UK MNDA DNA Bank ^65^ were originally derived from sixteen different individuals: 4 healthy individuals (without ALS), carrying the canonical IL18RAP 3’UTR sequence (Control; Canonical IL18RAP 3’UTR); 4 sporadic ALS patients, carrying the canonical IL18RAP 3’UTR sequence (sALS; Canonical IL18RAP 3’UTR); two healthy individuals, carrying a variant form of IL18RAP 3’UTR (Control; Variant IL18RAP 3’UTR); two sporadic ALS patients carrying a variant form of IL18RAP 3’UTR (sALS; Variant IL18RAP 3’UTR) and 4 C9orf72 ALS patients, carrying the canonical IL18RAP 3’UTR sequence (C9orf72; Canonical IL18RAP 3’UTR) (Cell lines listed in Supplementary Table 8; Weizmann IRB: 537-1). LCLs were cultured in RPMI-1640 (Gibco, 21875091) with 20% inactivated fetal bovine serum (FBS, Biological Industries, 04-001-1A), 1% L-glutamine and 1% penicillin-streptomycin (Biological Industries, 03-0311B) at 37°C, 5% CO2. Human Bone Osteosarcoma Epithelial Cells (U2OS), were maintained in Dulbecco’s Modified Eagle Medium (DMEM, Biological Industries, 01-050-1A) supplemented with 10% FBS, 1% penicillin-streptomycin at 37°C, 5% CO2. Human iPSCs were cultured on Matrigel (Corning, 354277) coated plated in mTeSR1 medium (Stemcell technologies, 85850) according to the manufacturer’s instructions. Briefly, cells were passaged at 70–90% confluent with StemPro accutase (Gibco, A11105-01) and seeded in mTeSR1 medium supplemented with 10 nM Y-27632 dihydrochloride (Tocris, 1254). Cells were refreshed with mTeSR1 medium every 24 hours until passaged.

### Isolation and Culture of Rat Cortical Astrocytes

All experiments were performed in accordance with relevant guidelines and regulations of the Institutional Animal Care and Use Committee at Weizmann Institute of Science (IACUC 09491120-1). Primary cortical a**s**trocytes were isolated and cultured as previously described ^96^ with several modifications. Briefly, the cerebral cortex of postnatal day 1 (P1) Sprague-Dawley rat pups was dissected and placed in DMEM/F12 containing 0.5% trypsin (biological industries, 03-046-5B). After 30 min incubation at 37 °C water bath, the cortical tissues were mechanically dissociated with pipette into single cells and were seeded on poly-D-lysine (Sigma Aldrich, 7405) coated T75 culture flask in Astrocytes medium (DMEM/F12 (Gibco, 31330) supplemented with 10% FBS, 50U/mL Pen-strep and 2Mm Glutamax (Gibco, 35050-038)). The confluent cultures were shaken for 4 hours at 200 rpm to remove microglial cells. Each T75 flask was trypsinized and splited into three new T75 flasks. After 7-8 days the confluent flasks were trypsinized and were frozen (in 90% FBS, 10% DMSO) until further use.

### I^3^LMNs neuronal differentiation and *Syn::GFP+* transduction

Differentiation of hiPSCs into lower motor neurons (i^3^LMNs, iPSCs containing Doxycycline induced human NGN2, ISL1, and LHX3 (hNIL)) was performed as described previously ^76^. Briefly, iPSCs were seeded on day 0 into mTeSR1 medium supplemented with 10 nM Y-27632 dihydrochloride. Few hours after seeding, cells were transduced with *Syn::GFP* lentivirus (pHR-hSyn-EGFP, Addgene #114215). 24 hrs. after seeding the cells medium was replaced with differentiation medium (DMEM/F12 (Gibco, 31330-038) with 1× MEM non-essential amino acids (Gibco, 11140-035), 2mM GlutaMAX (Gibco, 35050-038), 1× N-2 supplement (Gibco, 17502-048), 2 μg/mL Doxycycline (Sigma Aldrich, D9891-1G.) and 10 nM Y-27632 dihydrochloride). On day 3, cells were split using accutase, counted and re-seeded on poly-D-Lysine coated dishes containing Rat astrocytes in neuronal medium (B27 Electrophysiology medium (Gibco, A14137-01) supplemented with 1× MEM non-essential amino acids, 2mM GlutaMAX, 1× N-2 supplement, and 1 μg/mL mouse laminin (Gibco, 23017-015)). Twice a week half of the media was removed, and an equal volume of fresh media was added.

### Generation of IL18RAP 3’UTR rare variant hiPSCs lines

iPSCs were generated by the Ichida lab from human lymphocytes from ALS patients obtained from the National Institute of Neurological Disorders and Stroke (NINDS) Biorepository at the Coriell Institute for Medical Research. lymphocytes were reprogrammed into iPSCs as previously described ^66^. The NINDS Biorepository requires informed consent from patients.

Human iPSC lines were maintained on irradiated MEFs in hESCs medium [DMEM/F12 (Sigma-Aldrich, D6421) supplemented with 20% KO Serum Replacement (Gibco, 10828-028,), 1% GlutaMax (Gibco, 35050038), 1% MEM-NEAA (Biological Industries, 01-040-1A), 0.1mM 2-Mercaptoethanol (31350-010, Gibco), 10ng/ml hFGF (PeproTech, 100-18B)] and passaged twice a week with Collagenase IV (Worthington, LS004188).

CRISPR guides were chosen using several design tools, including: the MIT CRISPR design tool ^97^ and sgRNA Designer, Rule set 2 ^98^, in the Benchling implementations (www.benchling.com), SSC ^99^, and sgRNAscorer ^100^, in their websites.

Prior to CRISPR procedure iPSCs were passaged once in feeder-free condition [LDEV Free GelTrex matrix (Gibco, A1413202), mTESR1 medium (StemCell Technologies, 85850)], dislodged as single cells using StemPro Accutase (Gibco, A11105-01), washed twice with Opti-MEM (Gibco, 31985-047) and counted. 90ul cells suspension containing 1M cells was mixed with 10 uL DNA mix: 4 ug pSpCas9(BB)-2A-Puro (PX459) plasmid (Addgene #48139), • 0.4 ug gRNA encoding plasmid (pKLV-U6gRNA(BbsI)-PGKzeo2ABFP, derived from pKLV-U6gRNA(BbsI)-PGKpuro2ABFP (Addgene)), 1 ug (8 pmol) ssODN repair template (Supplementary Table 15) (IDT, 400 bases Megamer DNA Oligonucleotide) and 2.6 ug carrier plasmid DNA. CRISPR reaction components were introduced to iPSCs by single round electroporation using Nepa21 system (NEPA GENE). 100 uL cells and DNA suspension was transferred to Nepa Electroporation Cuvette 2 mm gap (Nepa Gene, EC-002). Electroporation conditions: 150 V Poring pulse; 5 ms Pulse length; 20 V Transfer pulse; 50 ms Pulse length. Electroporated cells were transferred to two GelTrex coated 100 mm dishes (1K and 10K) in mTeSR medium supplemented with 10 uM ROCK inhibitor (PeproTech, 1293823) and placed into CO2 incubator for 2 days. 48h past electroporation cells were treated with • 0.5 ug/mL Puromycin (Sigma-Aldrich) for 2 consecutive days. Survived cells were maintained until clones development. Single clones were picked and transferred to 96 well plates. Matured clones were genotyped at the first passage. Additionally, the top five predicted off-target sites for the guide RNA were sequenced (Supplementary Table 16). Selected clones containing desired mutations were expanded, cryopreserved, and used for the downstream experiments.

### Differentiation and culturing of human iPSC-derived microglia

hiPSCs were differentiated to microglia-like cells as previously described ^68^. Briefly, to form embryoid bodies (EBs), iPSCs were seeded into 96 well suspension plates in mTeSR1 media supplemented with 50 ng/mL rhBMP4 (Peprotech, 314-BP), 50 ng/mL VEGF (Peprotech, 100-20), 20 ng/mL SCF (Peprotech, 300-07) and 10 nM Y-27632 dihydrochloride. Everyday half of the medium was removed, and an equal volume of fresh media was added. After four days 12 EBs were transferred into each well of 6 well plate in X-VIVO 15 (Lonza, BE02-060Q) containing 100 ng/mL M-CSF (Peprotech, 300-25), 25 ng/mL IL-3 (Peprotech, 200-03), 2 mM Glutamax, 55 uM 2-mercaptoethanol (Gibco, 31350-10) and 100 U/mL penicillin/streptomycin (Biological Industries, 03-031-1B). iPSC derived progenitor microglia (ipMG) were collected weekly from the supernatant and were co-cultured with iPSC derived neurons in 96 well plates (Greiner, 655090) in neuronal medium containing 10 ng/mL IL-34 (Peprotech,200-34). EB medium was refreshed weekly.

### i^3^LMNs survival assay

Survival assay was conducted by monitoring eGFP signal of day 5 i^3^LMNs co-cultured with two independent CRISPR-edited isogenic iPSC-derived microglia (harboring canonical or variant IL18RAP 3’UTR), with C9orf72 genetic background. Cells were monitored for over 20 days using Incucyte® Live-Cell Analysis System (Sartorius). Daily longitudinal microscopic tracking was performed following Lipopolysaccharide (LPS, 100 ng/mL) and IL-18 treatment (100 ng/mL). i^3^LMNs survival assay was performed using three individual replicates for each line, with 3-8 co-culture wells per condition. Twice a week half of the media was removed, and an equal volume of fresh media containing LPS and IL-18 was added.

### Cloning

Full IL18RAP coding sequence (CDS) and 3’UTR sequence (2223bp) in pMX vector was purchased from GeneArt (Invitrogen, Supplementary Table 15) and subcloned with V5 epitope into pcDNA3. Different mutants, including: WT IL18RAP CDS + mutant 3’UTR (V1 or V3), and a dominant-negative coding mutant E210A-Y212A-Y214A CDS + WT 3’UTR (3CDS) ^41^ created by Transfer-PCR mutagenesis^101^. Next, WT and mutants full IL18RAP were subcloned into pUltra vector (a gift from Malcolm Moore, Addgene plasmid #24130, for which mCherry was replaced with EGFP), downstream of the human Ubiquitin C promoter and EGFP-P2A. Cloning procedures were done via restriction-free cloning ^102^. List of primers used for cloning and Transfer-PCR mutagenesis described in Supplementary Table 16.

### Transfection

Transfection to U2OS cells at 1.9 cm^2^ corning plates was performed at 70–80% confluence, 24 hrs. post-plating in antibiotic-free media, using Lipofectamine 2000, 0.5 µL per well (Thermo Fisher Scientific, Cat# 11668027). Each well was considered as a single replicate. For NF-κB reporter assay, U2OS cells were induced with/without recombinant IL-18 (5 ng/mL) 72 hrs. post-transfection with full coding sequence of IL18RAP coding region + 3’UTRs (pUltra vector 500 ng / 1.9 cm^2^ plate), luc2P/NF-κB-RE (pGL4.32 100 ng) luciferase and Renilla luciferase (hRluc 10 ng). Following 6 hrs. cells were harvested for Dual-Luciferase® Reporter Assay (E1960) and luminescence was quantified using Veritas™ Microplate Luminometer.

### RNA extraction, cDNA synthesis, and quantitative real-time PCR

Total RNA from LCLs was extracted using Direct-Zol RNA MiniPrep (Zymo Research ,R2052) according to manufacturer instructions. Total RNA from ipMGs was extracted using miRNeasy micro Kit (QIAGEN, 217084) according to manufacturer instructions. Total RNA was reverse transcribed using High Capacity cDNA Reverse Transcription Kit (applied biosystem, 4368814) according to manufacturer instructions, except for the mRNA stability assay, where equal volume of RNA (and not equal amounts of RNA) from each sample was used to generate cDNA. Quantitative Real-time PCR was performed using TaqMan Universal PCR master Mix (applied biosystem, 4304437) or KAPA SYBR FAST (Roche, KK4605). Primers and TaqMan probes are shown in Supplementary Table 16.

### Bulk MARS-Seq

200,000 ipMGs harboring variant or canonical IL18RAP 3’UTR (n=4) were treated with 100 ng/mL LPS + 100 ng/mL IL-18 for 6 hrs. in ipMG media (Advanced DMEM (Gibco, 12491-015) containing 1× N-2 supplement (Gibco, 17502-048), 2mM GlutaMAX (Gibco, 35050-038), 55 uM 2-mercaptoethanol (Gibco, 31350-10), 50 U/mL penicillin/streptomycin (Biological Industries, 03-031-1B) and 100 ng/mL IL-34 (Peprotech, 200-34). Following 6 hrs. RNA was extracted as described above and a bulk adaptation of the MARS-Seq protocol (Jaitin et al., Science 2014; Keren-Shaul et al., Nature Protocols, 2019) was used to generate 3’ RNA-Seq libraries for expression profiling. Briefly, 50 ng of input RNA from each sample was barcoded during reverse transcription and pooled. Following Agencourt Ampure XP beads cleanup (Beckman Coulter), the pooled samples underwent second strand synthesis and were linearly amplified by T7 in-vitro transcription. The resulting RNA was fragmented and converted into a sequencing-ready library by tagging the samples with Illumina sequences during ligation, RT, and PCR. Libraries were quantified by Qubit and TapeStation as well as by qPCR for GAPDH gene as previously described (Jaitin et al., Science 2014; Keren-Shaul et al., Nature Protocols, 2019). Sequencing was done on a NovaSeq 6000 system, SP Reagent Kit, 100 cycles (Illumina; paired-end sequencing).

Analysis of the MARS-seq was done using the UTAP pipeline (^103^; the Weizmann Institute Bioinformatics Unit) to map the reads to the human genome and to calculate Unique Molecule Identifier (UMI) counts per gene. Reads were trimmed from their adapter using cutadapt (parameters: -a AGATCGGAAGAGCACACGTCTGAACTCCAGTCAC -a “A(10)” –times 2 -u 3 -u −3 -q 20 -m 25) and mapped to hg38 genome (STAR v2.4.2a). The pipeline removes UMI redundancy and quantifies the 3’ of RefSeq annotated genes (1,000 bases upstream and 100 bases downstream of the 3’ end). Genes having a minimum of five reads in at least one sample were considered for further analysis. Differentially expressed (DE) gene detection and count normalization analysis were performed by DESeq2. P-values in the UTAP results were adjusted for multiple testing using the Benjamini and Hochberg procedure. Thresholds for significant DE genes: padj < 0.01, |log2FoldChange| >= 0.585, baseMean > 20. This assay was done with critical advice from Dr. Hadas Keren-Shaul from the Genomics Sandbox unit at the Life Science Core Facility of Weizmann Institute of Science.

### Cell lysis and Western blot

LCLs were washed in PBSx1, centrifuged at 800 × *g* for 5 min at 4°C, pelleted, and lysed in ice-cold RIPA buffer (Supplementary Table 17) supplemented with cOmplete™ Protease Inhibitor Cocktail (Roche, 4693116001) and PhosSTOP™ (Roche, 4906837001). The lysates were cleared by centrifugation at 15,000 × g for 10 min at 4°C. Protein concentrations quantified with Protein Assay Dye Reagent (Bio-Rad, 500-0006), resolved at 30-50μg of total protein/well by 8-10% polyacrylamide / SDS gel electrophoresis at 100-120 V for 70 min. After gel electrophoresis proteins were transferred to nitrocellulose membrane (Whatmann, 10401383) at 250 mA for 70 min. Membranes were stained with Ponceau (Sigma, P7170), blocked for 1 hour at RT with 3% Bovine albumin fraction V (MPBio 160069) or 5% milk protein in PBST (PBS containing 0.05% TWEEN-20), and then incubated with primary antibodies (see Supplementary Table 18) O.N. at 4°C with rocking in antibody-Solution [5% albumin, 0.02% sodium azide, 5 drops of phenol red in 0.05% PBST]. Following primary antibody incubation, membranes were washed 3 times for 5 min at RT with 0.05% PBST then incubated for 1 hour at RT with horseradish peroxidase (HRP)-conjugated species-specific secondary antibodies, washed 3 x 5 min in 0.05% PBST at RT, and visualized using EZ-ECL Chemiluminescence (Biological Industries, 20500-120) by ImageQuant™ LAS 4000 (GE Healthcare Life Sciences). Densitometric analysis was performed using ImageJ (NIH).

### In-Vitro Transcription of biotinylated IL18RAP 3’UTR

To identify the potential trans-acting factors that might differentially bind to the canonical and variant 3’UTRs an RNA-pulldown and mass spectrometry assay was performed on *in vitro* transcribed canonical and variant forms of the IL18RAP 3’UTRs, V1 and V3. Briefly, The canonical, V1 and V3 biotinylated-IL18RAP 3’UTR sequences (384nt), and the negative control (ultrapure water only), were produced by using in vitro transcription HiScribe™ T7 ARCA Kit (NEB, E2060S) following the manufacturer instructions. Briefly, 300 ng of purchased DNA template (50 ng/uL) (Twist, Supplementary Table 15) was incubated with unlabeled ATP/GTP/CTP and 5% biotin-labelled UTP, at 37°C for 3 hrs. Next, DNase treatment was performed by incubating the reactions at 37°C for 30 min and was followed by incubation at 65°C for 10 min to terminate the reaction. The RNA products were purified by an RNA cleanup purification kit (Zymo Research, R1015). The concentrations of the purified RNA samples were measured by nanodrop and the expected length was analyzed by TapeStation.

### Pull Down of IL18RAP 3’UTR RNA-associated proteins

LCL cell pellets were suspended and lysed in RIPA buffer followed by centrifugation at 15,000xg for 10 min at 4°C. The concentrations of the cleared supernatants were measured by Bradford assay. 1 mg lysate per sample was incubated with Pierce streptavidin magnetic beads (Thermo Scientific, 88817) for 30 min at 4°C in rotation, to pre-clear the lysates from endogenous biotinylated-proteins. To bind IVT products (WT, V1, V3 and negative control; n=6 repeats/group) to the beads, new prepared binding Pierce streptavidin magnetic beads were incubated by rotation with equal amounts of IVT products for 30 min at 4°C (100 uL beads/10 pmol RNA product). After 30 min, the tubes of incubated IVT products with beads were washed three times, and then the cleared lysate was added equally to each tube and incubated for 30 min at 4°C. In the next step, the samples were washed three times by magnetizing the beads and resuspended by vortex with a high salt buffer. The bound beads were magnetized and suspended in 20 ul RNase-free PBSx1 for on-bead digestion procedure.

### Liquid Chromatography and Mass Spectrometry

The resulting peptides were analyzed using nanoflow liquid chromatography (nanoAcquity) coupled to high resolution, high mass accuracy mass spectrometry (Q-Exactive HF). Each sample was analyzed on the instrument separately in a random order in discovery mode.

### Raw proteomic data processing

Raw MS data were processed using MaxQuant version 1.6.6.0 (Cox and Mann, 2008). Database search was performed with the Andromeda search engine (Cox and Mann, 2011; Cox et al., 2011) using the human Uniprot database, appended with common lab protein contaminants. Forward/decoy approach was used to determine the false discovery rate and filter the data with a threshold of 1% false discovery rate (FDR) for both the peptide-spectrum matches and the protein levels. The label-free quantification (LFQ) algorithm in MaxQuant (Cox et al., 2014) was used to compare between experimental samples. Additional settings included the following modifications: Fixed modification-cysteine carbamidomethylation. Variable modifications-methionine oxidation, asparagine and glutamine deamidation, and protein N-terminal acetylation.

### Proteomics statistical analysis

ProteinGroups output table was imported from MaxQuant to Perseus v.1.6.2.3 environment (Tyanova et al., 2016). Quality control excluded reverse proteins, proteins identified only based on a modified peptide, and contaminants. Non-specific streptavidin-bead binders were excluded by the following procedure: LFQ Intensity values were log2 transformed, and two outlier samples were excluded from further analysis. Missing values were imputed by creating an artificial normal distribution with a downshift of 1.8 standard deviations and a width of 0.4 of the original ratio distributions. Student’s t-test with S0 = 0.1 was performed with FDR P-value ≤ 0.05 between the experimental groups (Canonical, V1 and V3) and the negative control group, which was defined as a single control group. Proteins that passed all QC filters were separated for each of the experimental groups and compared to the negative control samples (ultrapure water). The statistically significant-associated proteins were filtered to retain only proteins that were found in 50% of the repeats in at least one experimental group and were represented by at least one unique peptide. The enriched proteins were subjected to student’s t test between every two groups (canonical vs. V1 and canonical vs. V3), with S0 = 0.1, FDR P-value ≤ 0.05 and fold-change threshold >2.

### Processing of Mouse Brain Samples for Flow Cytometry

Wild-type C57BL/6 mice were euthanized with CO2 and perfused with PBS through the left ventricle of the heart. Dissected mouse cortex was cut into smaller pieces using scissors and digested in 0.5 mg/mL Collagenase IV (Worthington Biochemical), 10 µg Deoxyribonuclease (Sigma-Aldrich), 10% HI-FBS, RPMI1640 (Gibco) at 37°C for 30 minutes with continuous agitation. Digested samples were gently triturated for 1 minute and the enzymatic reaction was stopped by adding 1 mM EDTA in PBS. The homogenate was filtered through a 100 µm cell strainer and centrifuged at 400 x g for 8 minutes at 4°C to pellet the cells and myelin. This was followed by myelin removal step by gradient centrifugation with 30% Percoll (Sigma-Aldrich) in PBS (700 x g for 20 minutes at 21oC; without brakes during deceleration). After myelin (the top white layer) separation, the middle transparent layer was collected, washed in PBS, and centrifuged at 400 x g for 8 minutes at 4°C to pellet the cells.

Cells pellets were incubated with Mouse Fc block (BD Biosciences 553142), Fixable Viability Stain 620 (BD Biosciences 564996) and the following antibody mixture in PBS at 4°C for 30 minutes: BV421 Rat Anti-CD11b (BD Biosciences 562605), BV510 Hamster Anti-Mouse TCR β Chain (BD Biosciences 563221), BV711 Rat Anti-Mouse Ly-6G (BD Biosciences 563979), APC-Cy7 Rat Anti-Mouse CD45 (BD Biosciences 557659), and Polyclonal Goat IgG Anti-Mouse IL-18Rβ (R&D Systems AF199). Samples were then washed with PBS and incubated with Alexa Fluor 647 Donkey Anti-Goat IgG (H+L) Cross-Adsorbed Secondary Antibody (Invitrogen A-21447) in PBS at 4°C for 30 minutes. Surface-stained samples were washed with PBS and fixed and permeabilized with BD Fixation/Permeabilization solution (BD Biosciences 554714) at 4°C for 30 minutes, followed by intracellular staining with Alexa Fluor 488 Anti-NeuN Antibody (EMD Millipore MAB377X) and eFluor 570 Anti-GFAP (eBioscience 41-9892-82) in BD Perm/Wash Buffer (BD Biosciences 554714) at 4°C for 30 minutes. Cells were washed with BD Perm/Wash Buffer and resuspended in PBS for analysis with a FACSymphony (BD Biosciences). Data were collected as FCS files and analyzed with FlowJo v10 software (BD Biosciences). Antibody specificity was assessed using relevant isotype control antibodies and fluorescence minus one. Compensation was adjusted using single-stained samples.

The expression of IL-18RAP (IL-18Rβ) was expressed as Mean Fluorescence Intensity (MFI) or % frequency after gating for the following cell types: immune cells (CD45^hi^), microglia (MG: CD45^int^ CD11^hi^), neurons (CD45^-^ CD11b^-^ NeuN^+^), and astrocytes (CD45^-^ CD11b^-^ GFAP^+^). Animal procedures were approved by the Walter and Eliza Hall Institute Animal Ethics Committee (Ethics application: 2020.017).

### Statistical analysis

Statistics performed with Prism Origin (GraphPad). Shapiro-Wilk test was used to assess normality of the data. Pairwise comparisons passing normality test were analyzed with Student’s *t*-test, whereas the Mann-Whitney test was used for pairwise comparison of nonparametric data. Multiple group comparisons were analyzed using ANOVA with post hoc tests. For age of diagnosis and age of death a Permutation Test was used (a Monte-Carlo simulation test on the t-test between ALS patients harboring canonical or variants of the IL18RAP 3’UTR). Statistical P-values <0.05 were considered significant. Data are shown as scatter dot plot with mean and SEM, box plots, or as noted in the text.

## Supplementary Materials

Fig. S1. Study design.

Fig. S2. Region-based rare-variant association analyses.

Fig. S3. 3′UTR-based rare-variant association analysis, using different algorithms, and illustration of rare variants identified in the IL18RAP 3′UTR.

Fig. S4. Restricting burden analysis to the proximal part of 3’UTRs does not improve the association signal.

Fig. S5. IL18RAP and p-NF-κB expression is elevated in lymphoblastoid cells from patients with the C9orf72 repeat expansion.

Fig. S6. IL18RAP 3’UTR variant attenuates IL-18 - NF-κB signaling in U2OS cells.

Fig. S7. IL18RAP is mainly expressed on mouse microglia cells.

Fig. S8. Evaluation of IL18RAP and IL-18 mRNA expression in motor neurons of patients with ALS.

Fig. S9. iPSC-derived microglia express the microglial-specific marker, TMEM119.

Fig. S10. Differentially bound RNA binding proteins to variant 3’UTR (V3) relative to canonical 3’UTR.

Table S1. Total number of samples before and after quality control procedures, stratified by country.

Table S2. Samples quality control procedures.

Table S3. Candidate genes list.

Table S4. Number of rare genetic variants identified.

Table S5. Detailed description of variants in protein-coding sequences of *NEK1* and *SOD1* and the IL18RAP 3′UTR, in Project MinE discovery cohort.

Table S6. Identified IL18RAP 3′UTR variants in Project MinE discovery cohort.

Table S7. Identified IL18RAP 3′UTR variants in discovery and replication cohorts.

Table S8. Control and patient human cell lines information.

Table S9. Proteomics data from IL18RAP 3’UTR pull-down experiments.

Table S10. Gene set enrichment analysis of differentially bound proteins in canonical vs. V1 IL18RAP 3’UTR.

Table S11. Phenotypic and variant information for patients with protective 3’UTR variants in Project MinE and NYGC cohorts.

Table S12. MARS-seq data for mRNAs measured in isogenic ipMG harboring variant vs. canonical IL18RAP 3’UTR.

Table S13. KEGG Pathway and Gene Ontology (GO) Biological Process Enrichment Results.

Table S14. BED file containing genomic coordinates of regions of interest.

Table S15. IL18RAP oligonucleotides.

Table S16. List of primers.

Table S17. Medium formulation.

Table S18. Materials and Antibodies.

Data File S1. Source data for IL18RAP and p-NF-κB western blot studies in LCLs (Figure 3D).

Data File S2. Source data for IL18RAP and p-NF-κB western blot studies in Control vs. C9orf72 LCLs (Supplementary figure 5D).

Data File S3. Source data for IL18RAP western blot studies in isogenic microglia (Figure 4B).

Data File S4. Source data for motor neuron survival assays (Figure 6B,C).

Data File S5. Source data for pNF-κB western blot studies in isogenic microglia, following microglia activation (Figure 7B).

Project MinE ALS Sequencing Consortium PI List

NYGC ALS Consortium PI List

Supplementary movie. Motor neuron survival was significantly improved in the presence of microglia harboring variant IL18RAP 3’UTR relative to canonical IL18RAP 3’UTR.

## Acknowledgments

We gratefully acknowledge the contributions of all participants and the investigators who provided biological samples and data for Project Mine ALS sequencing consortium, the New York Genome Center (NYGC) ALS Consortium, the Genome Aggregation Database (gnomAD) and Trans-Omics for Precision Medicine (TOPMed) of the National Heart, Lung, and Blood Institute (NHLBI, https://www.nhlbiwgs.org/topmed-banner-authorship). We thank Michael Ward (NINDS, NIH) for sharing human inducible i^3^LMN cells. Samples used in this research were in part obtained from the UK National DNA Bank for MND Research, funded by the MND Association and the Wellcome Trust. We acknowledge sample management undertaken by Biobanking Solutions funded by the Medical Research Council at the Centre for Integrated Genomic Medical Research, University of Manchester. The authors would like to thank the NINDS Biorepository at Coriell Institute for iPSC cell lines used in this study. We thank Bernardo Oldak and Prof. Jacob Hanna for microglia differentiation protocols, Dr. Noga Kozer and Dr. Haim Barr for assistance with live cell imaging, Dr. Alon Savidor and Dr. Yishai Levin for mass spectrometry, Dr. Merav Shmueli, Dr. Yifat Merbl and Dr. Ron Rotkof for advice and protocols. We thank LSE for language and scientific editing. Some illustrations were created with BioRender. Hornstein lab is supported by friends of Dr. Sydney Brenner. EH is Head of Andi and Larry Wolfe Center for Research on Neuroimmunology and Neuromodulation and incumbent of Ira & Gail Mondry Professorial chair.

## Author contributions

C.E. and A.Si. led the project; C.E. and A.Si. contributed to research conception, design and interpretations and wrote the manuscript with E.H.; C.E., E.B., T.O., K.R.V.E., S.L.P., M.M., S.M.K.F., N.Y., J.C.-K., K.P.K., R.A.A.V.D.S., W.S., A.A.K., A.I., A.Sh., A.R.J., E.C., D.R., O.W., R.H.B.J., P.J.S., P.V.D., L.H.v.d.B., H.P.P., E.S., A.A.-C. and J.H.V. collected samples, were involved in the sequence analysis pipeline, phenotyping, variant calling, provided expertise or were involved in the genetic association analysis of rare non-coding variants in human patients with ALS; S.-T.H. and J.K.I. provided stem cells and initial data; S.B.-D., E.A., G.B. and H.M.-K. were involved in the design, generation and validation of CRISPR-edited IL18RAP isogenic iPSCs; H.M.-K. and Y.M.D. performed the pull down experiments of IL18RAP 3’UTR RNA-associated proteins and analyzed the proteomic data; A.Si. and C.E. established human iPSC- derived microglia differentiation and culturing protocols, performed motor neuron survival experiments and interpreted data; A.Si., N.R. and C.E. performed molecular biology studies in LCLs and U2OS cell lines, including reporter assays, qPCR and protein quantification by western blots; E.Y. performed Bulk MARS-Seq experiment; C.-H.Y., C.L. and S.L.M. provided expertise and processed mouse cortex samples for Flow Cytometry; Y.C., Y.E.-A., S.W. and D.P.S. helped performing research; E.H. conceived and supervised the study and wrote the manuscript with C.E. and A.Si. All co-authors provided approval of the manuscript.

## Data availability

Human genetics data is publically available from the sequencing consortia: Project Mine ALS sequencing consortium, the New York Genome Center (NYGC) ALS Consortium, the Genome Aggregation Database (gnomAD), and NHLBI’s Trans-Omics for Precision Medicine (TOPMed). Gene Expression Omnibus accession number: GSE186757. All Other data used for this manuscript are available in the manuscript.

## Code availability

Variant annotation scripts are available at GitHub: https://github.com/TsviyaOlender/Non-coding-Variants-in-ALS-genes-.

---

**Supplementary Fig. 1.**
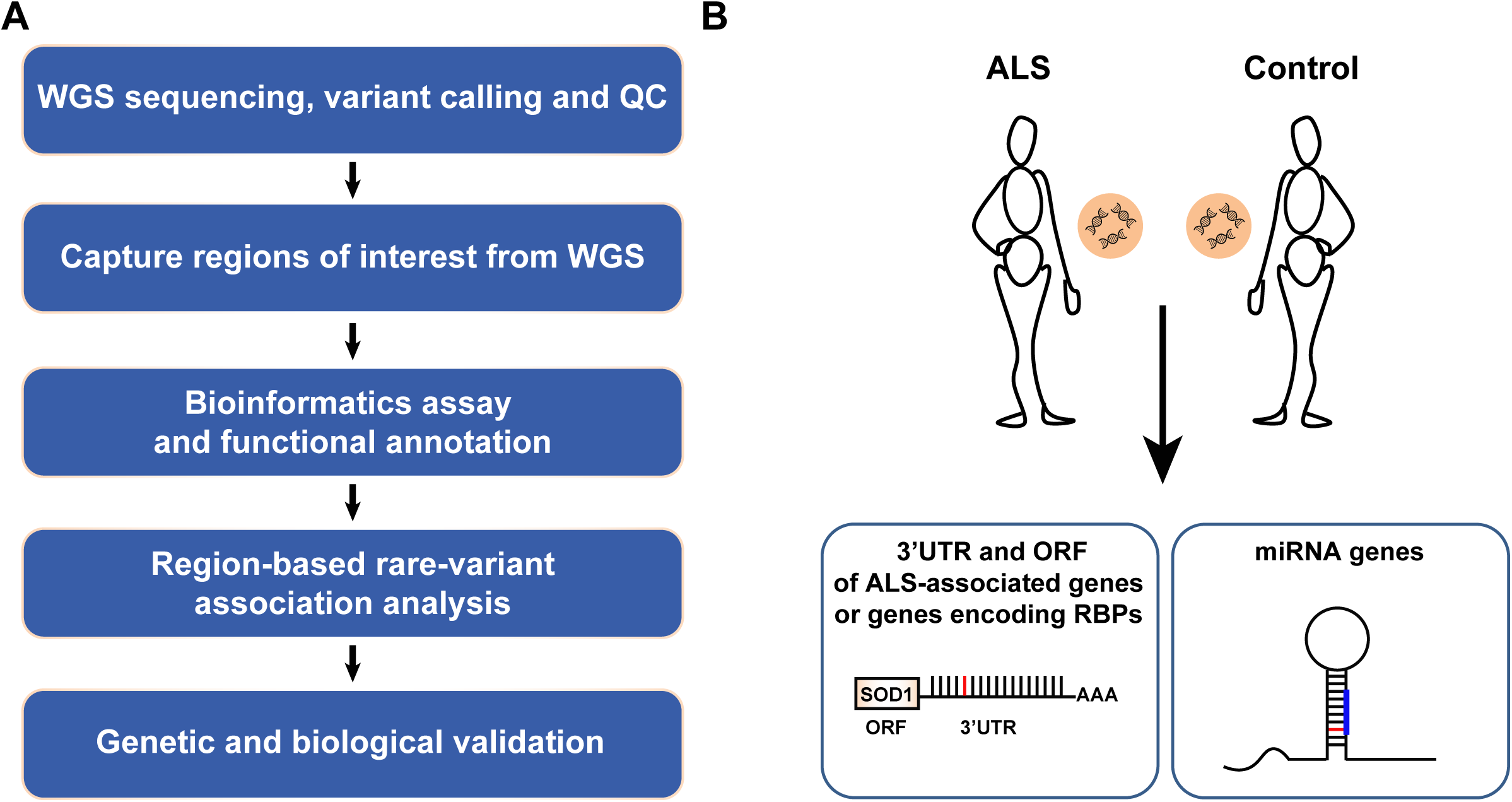
Study design. **(A)** Flow chart of approach for discovery of region-based rare-variants in non-coding genomic regions via association studies and **(B)** diagram depicting regions of interest comprising of 1,750 autosomal human pre-miRNA genes, 295 open reading frames encoding for proteins of interest, and 295 3′UTRs.

**Supplementary Fig. 2.**
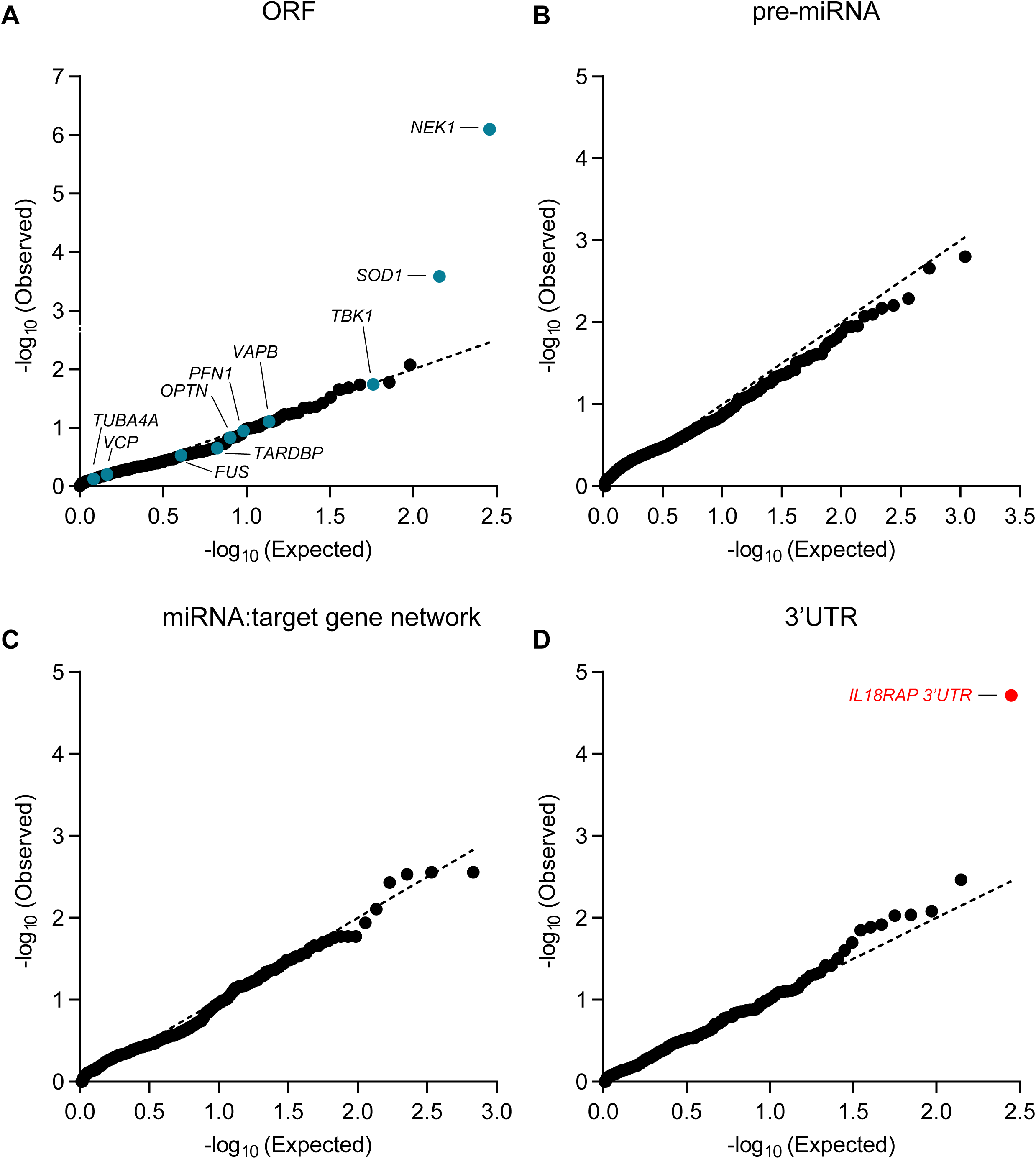
Region-based rare-variant association analyses. **(A-D)** QQ plot of obtained and expected P-values for the burden of rare-variants (log scale) gained by collapsed region-based association analysis of different genomic regions, comprised of (A) 295 candidate protein-coding regions listed in Supplementary Table 3. These ORFs encode for ALS-relevant proteins or proteins that are associated with miRNA biogenesis or activity. Variants were depicted if predicted to cause frameshifting, alternative splicing, abnormal stop codon or a deleterious non-synonymous amino acid substitution, in ≥ 3 of 7 independent dbNSFP prediction algorithms (genomic inflation λ = 0.96), (B) All known pre-miRNA genes in the human genome (genomic inflation λ = 1.31), (C) predicted networks, comprised of aggregated variants detected on a specific mature miRNA sequence and its cognate downstream 3’UTR targets (genomic inflation λ = 1.16), and (D) variants in 3′UTRs of the same 295 genes listed in Supplementary Table 3 (genomic inflation λ = 1.08). Data was obtained from 3,955 ALS cases and 1,819 controls (Project MinE). Features positioned on the diagonal line represent results obtained under the null hypothesis. Open-reading frames of 10 known ALS genes (blue). IL18RAP 3′UTR (red). P-values, calculated with SKAT-O.

**Supplementary Fig. 3.**
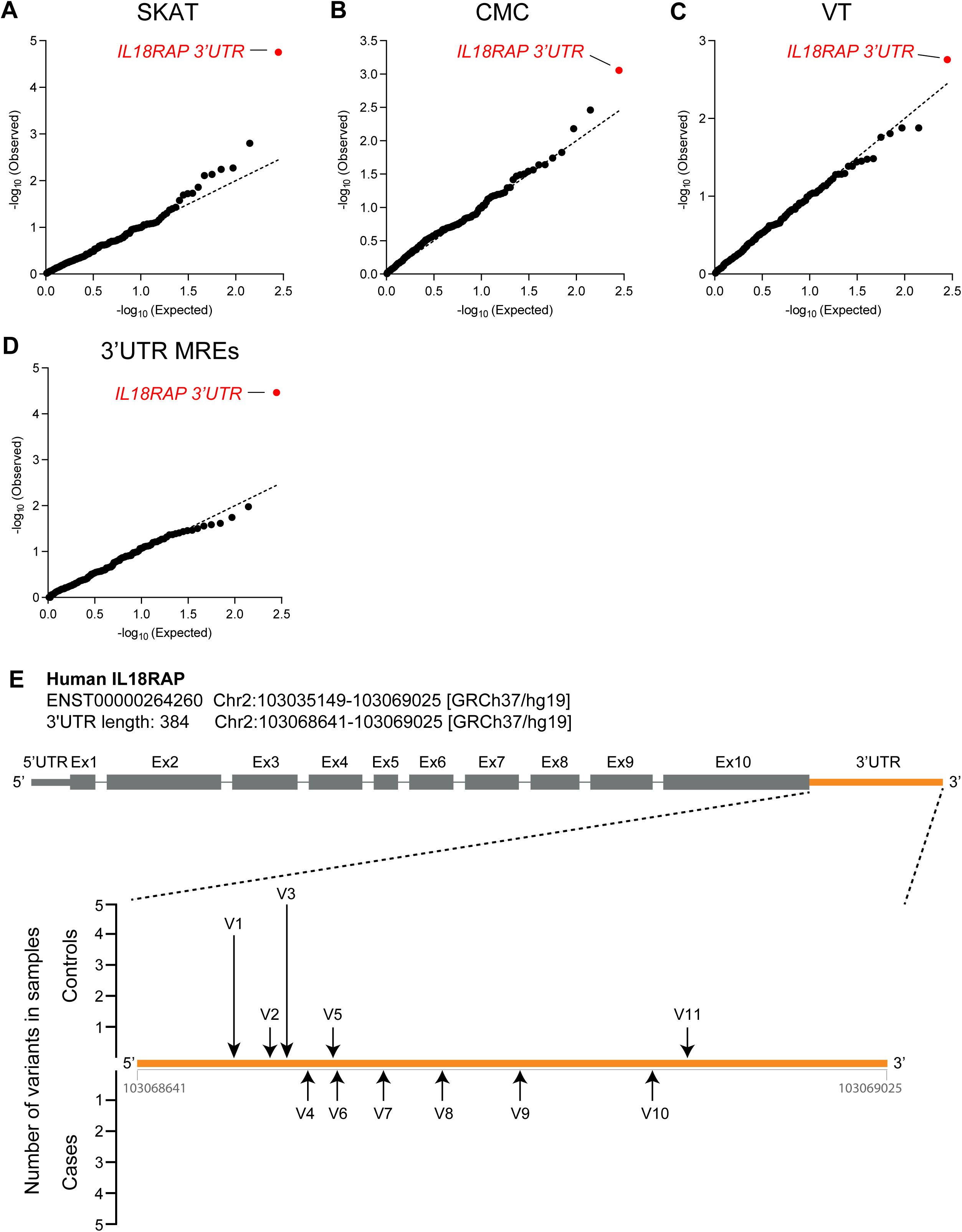
3′UTR-based rare-variant association analysis, using different algorithms, and illustration of rare variants identified in the IL18RAP 3′UTR. **(A-D)** QQ plot of obtained and expected P-values for the burden of rare variants (log scale) gained by collapsed region-based association analysis of genomic regions comprised of 295 3′UTRs listed in Supplementary Table 3, in the Project MinE cohort (3,955 ALS cases and 1,819 non-ALS controls). Features positioned on the diagonal line represent results obtained under the null hypothesis. IL18RAP 3′UTR (red) is the most significant 3’UTR associated with ALS using different algorithms: **(A)** Sequence Kernel Association Test, SKAT (genomic inflation λ = 1.02), **(B)** Combined Multivariate and Collapsing, CMC (genomic inflation λ = 1.34), **(C)** Variable Threshold with permutation analysis, VT (genomic inflation λ = 1.03). **(D)** IL18RAP 3′UTR also ranked as the top hit when aggregating variants abrogating or gaining miRNA recognition elements (MREs) in 3’UTRs (genomic inflation λ = 1.04). **(E)** Schematic of the IL18RAP transcript and 3′UTR (5’ to 3′) showing the number of control (upper) or ALS (lower) samples in which variants (black arrow) were identified in the Project MinE discovery cohort (Supplementary Table 6).

**Supplementary Fig. 4.**
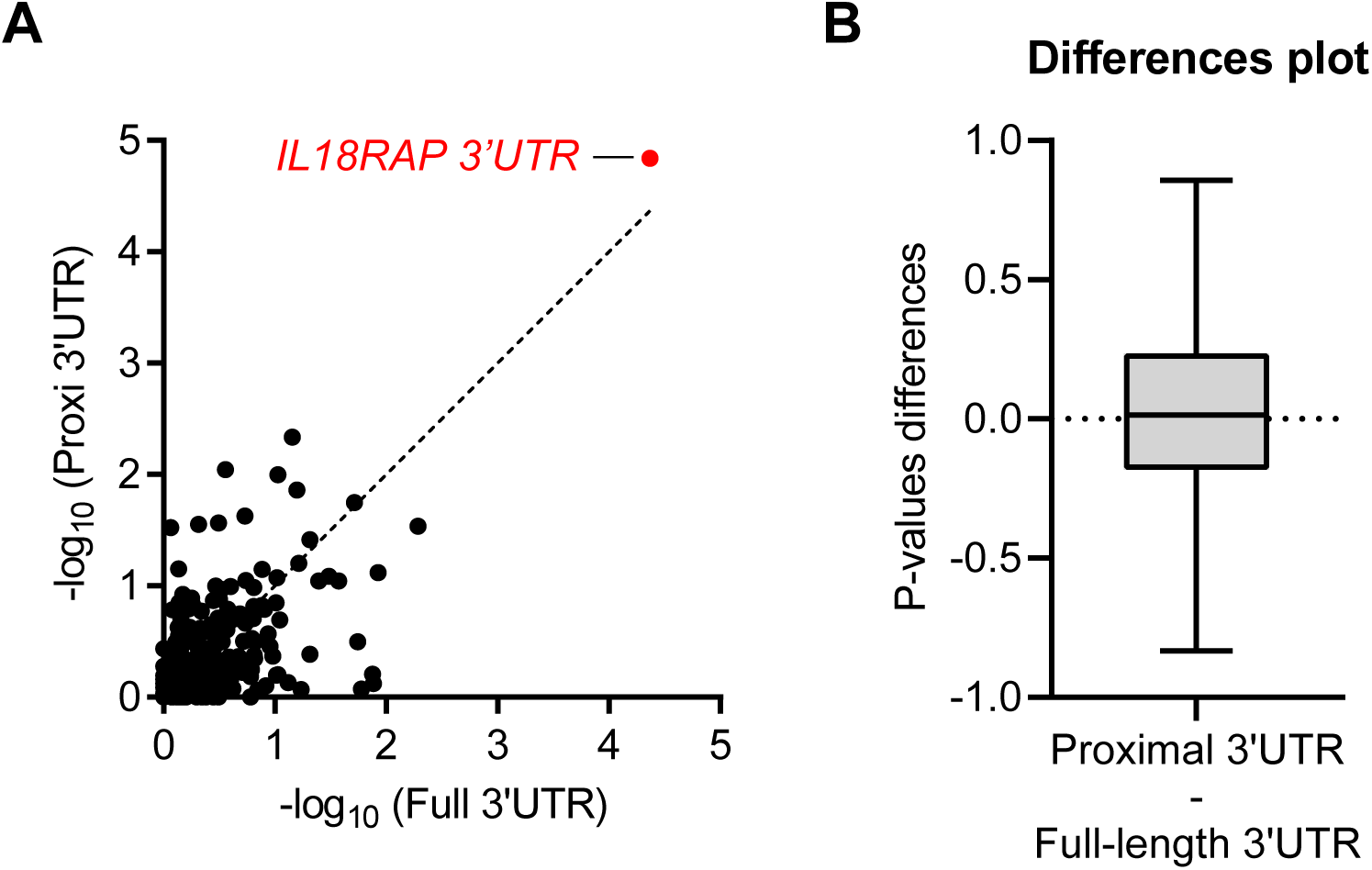
Restricting burden analysis to the proximal part of 3’UTRs does not improve the association signal. **(A)** Scatter plot with SKAT-O P-values (log scale) calculated for the burden of rare variants gained by collapsed region-based association analysis of the full 3’UTRs on the x-axis versus the 3’UTRs proximal quadrant on the y-axis, for the 295 3′UTRs listed in Supplementary Table 3, in the Project MinE cohort (3,955 ALS cases and 1,819 non-ALS controls) (Pearson correlation coefficient (r=0.61) and P-value ****<0.0001). The 45- degree diagonal line represents a perfect correlation of r=1. IL18RAP 3′UTR (red). **(B)** A Difference plot showing the difference between the two P-value measurements (3’UTRs proximal quadrant minus the full 3’UTRs). The bias (difference between means) is only 0.03. Overall the P-values gained from the 3’UTRs proximal quadrant were comparable to that of the full 3’UTRs in the cohort of 295 3’UTRs. Box plots depict median, upper and lower quartiles, and extreme points (Wilcoxon matched-pairs P-value > 0.05, Cohen’s d effect size = 0.1). Hence, the apparent spatial distribution of variants in IL18RAP 3’UTR seems to be a particular case, rather than part of a global pattern.

**Supplementary Fig. 5.**
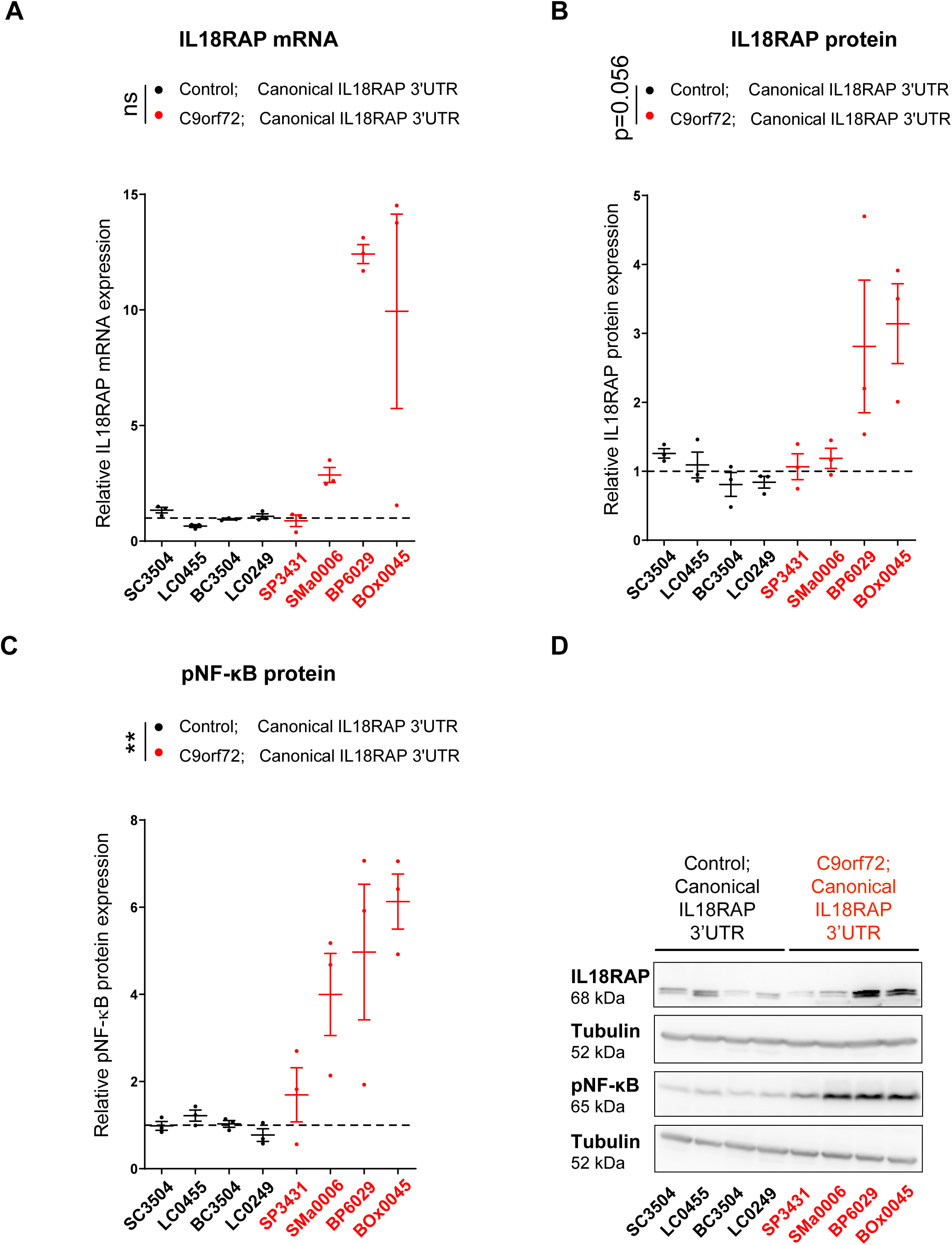
IL18RAP and p-NF-κB expression is elevated in lymphoblastoid cells from patients with the C9orf72 repeat expansion. **(A)** IL18RAP mRNA expression (qPCR normalized to IPO8 mRNA levels) and **(B)** IL18RAP or **(C)** p-NF-κB protein expression (Western blots, normalized to Tubulin). Extracts from eight different human lymphoblastoid cell lines (listed in Supplementary Table 8): Four lines of healthy individuals (without ALS) carrying the canonical IL18RAP 3’UTR sequence (Control; Canonical IL18RAP 3’UTR, black) and four C9orf72 ALS patients carrying the canonical IL18RAP 3’UTR sequence (C9orf72; Canonical IL18RAP 3’UTR, red). **(D)** Representative blots processed with anti-IL18RAP, anti p-NF-κB and anti-Tubulin antibodies. Mann-Whitney test (A) or one-sided student’s t-test with Welch’s correction on log-transformed data (B,C), was conducted based on the mean value of three independent passages for each of the eight human lymphoblastoid cell lines. Scatter dot plot with mean and SEM. ** P<0.01.

**Supplementary Fig. 6.**
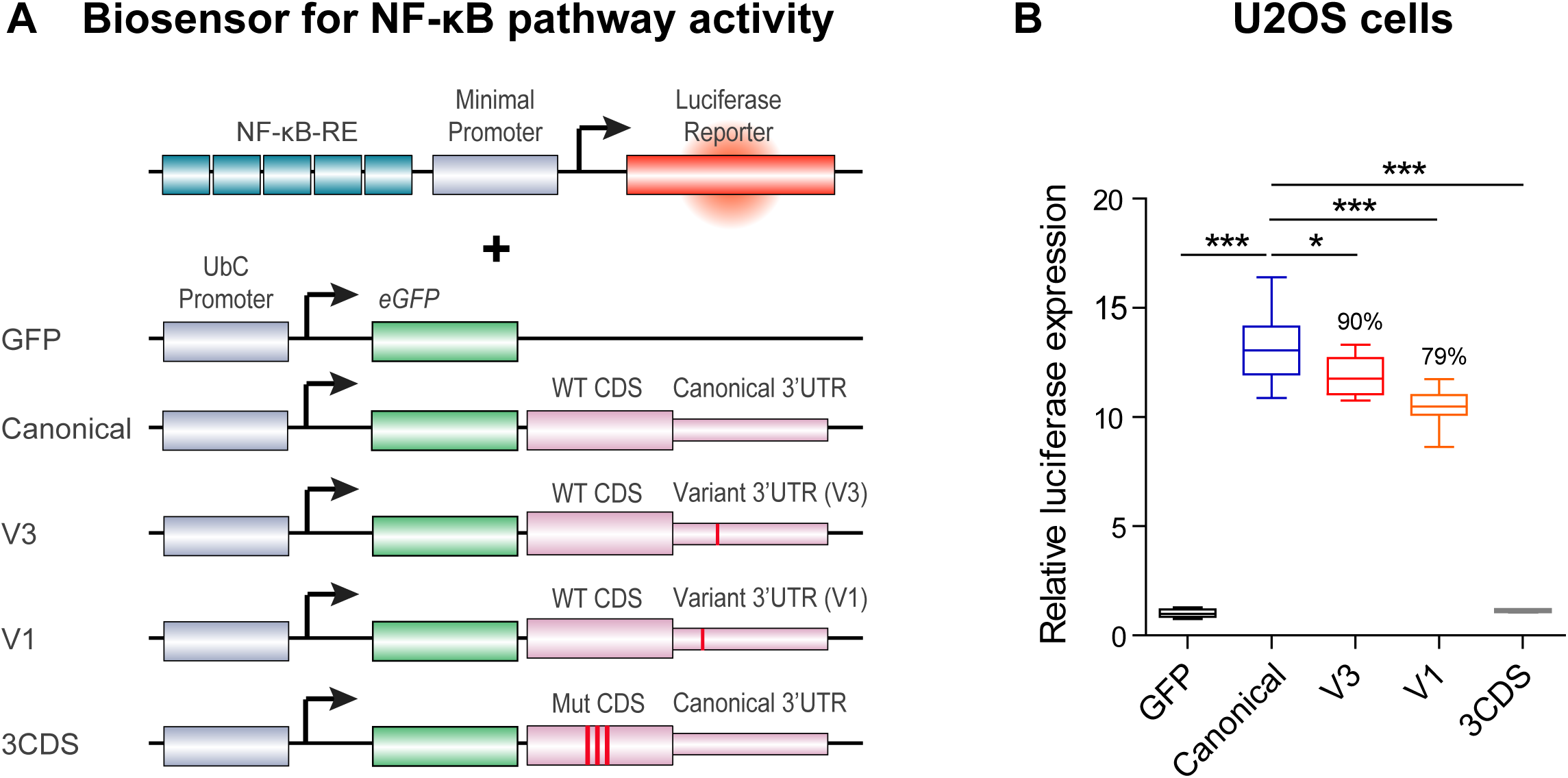
IL18RAP 3’UTR variant attenuates IL-18 - NF-κB signaling in U2OS cells. Diagram **(A)** and quantification **(B)** of NF-κB reporter assays in human U2OS cell line. To determine the ability of the IL18RAP variants V3 and V1 to induce NF-κB activity, U2OS cells were co-transfected with different IL18RAP coding region (CDS) and 3’UTR constructs (GFP, Canonical, V3, V1, n=9; 3CDS, *n*=4), along with an NF-κB activity reporter that drives luciferase (Luc2P) transcription via five copies of the NF-κB response element. NF-κB signaling was induced by adding human recombinant IL-18 to the medium. Variants V3 and V1 of the IL18RAP 3’UTR reduced NF-κB activity by ∼10% and ∼21%, respectively, relative to the WT IL18RAP 3’UTR. GFP vector and a dominant-negative coding mutant E210A-Y212A-Y214A CDS + WT 3’UTR (3CDS) ^41^, served as controls. Luciferase expression was normalized to transfected U2OS cells that were not induced with human recombinant IL-18. One-way ANOVA followed by Dunnett’s multiple comparison test was performed on square root-transformed data. Box plots depict median, upper and lower quartiles, and extreme points. * P<0.05; *** P<0.001. The experiment was repeated independently three times with similar results.

**Supplementary Fig. 7.**
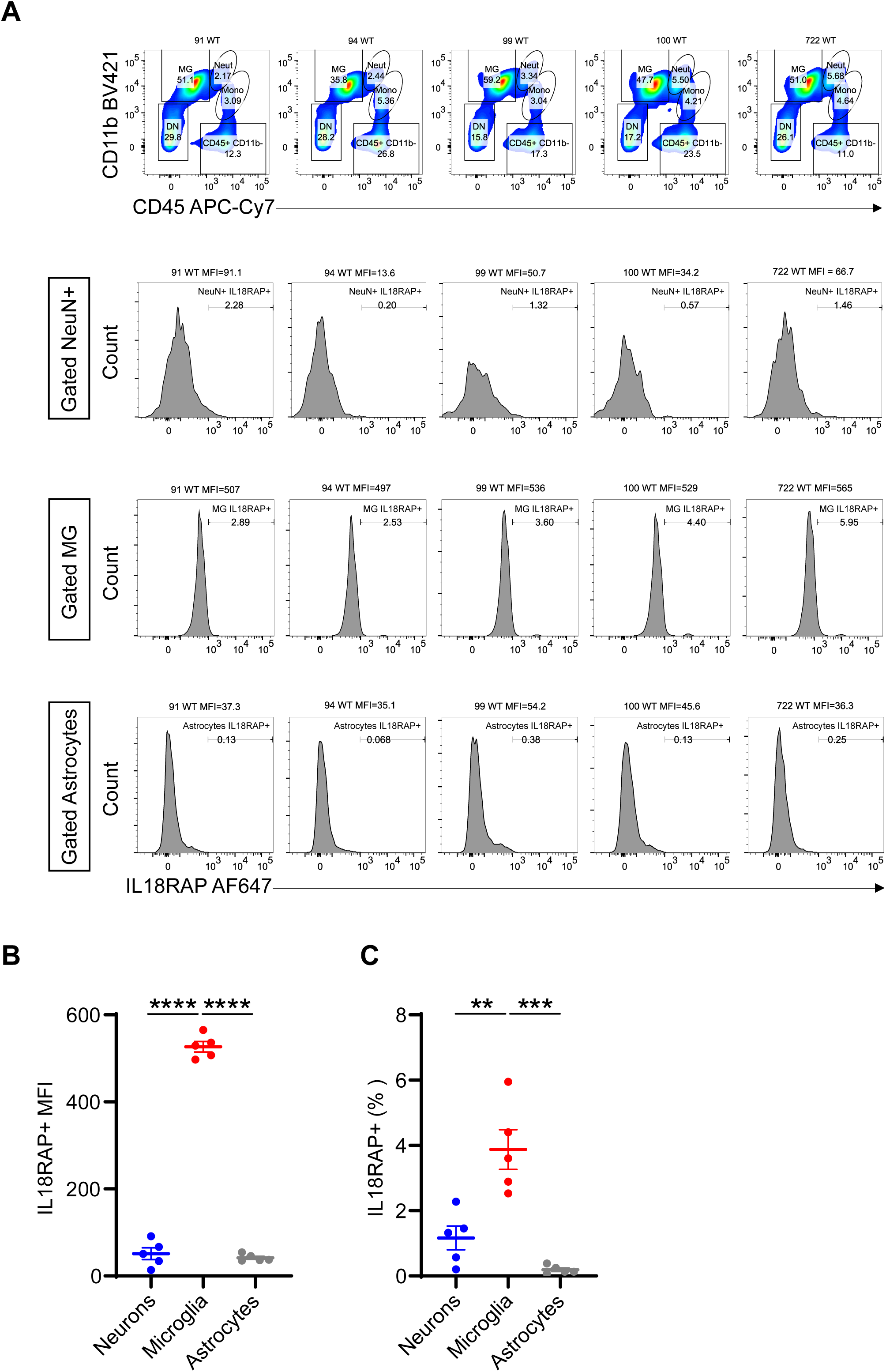
IL18RAP is mainly expressed on mouse microglia cells. (A-C) Flow cytometry was used to characterize IL18RAP expression levels in dissociated wild-type mouse cortex cells. The expression of IL-18RAP (IL-18Rβ) was expressed as Mean Fluorescence Intensity (MFI) and % frequency after gating for the following cell types: immune cells (CD45hi), microglia (MG: CD45int CD11hi), neurons (CD45-CD11b-NeuN+), and astrocytes (CD45- CD11b- GFAP+). FACS analysis reveals that IL18RAP is mainly expressed on microglia cells. A scatter dot plot with mean and SEM values for the median fluorescence intensity (MFI) and percentage of IL18RAP+ cells is shown. One-way ANOVA followed by Tukey’s multiple comparison test. ** P<0.01, *** P<0.001, **** P<0.0001.

**Supplementary Fig. 8.**
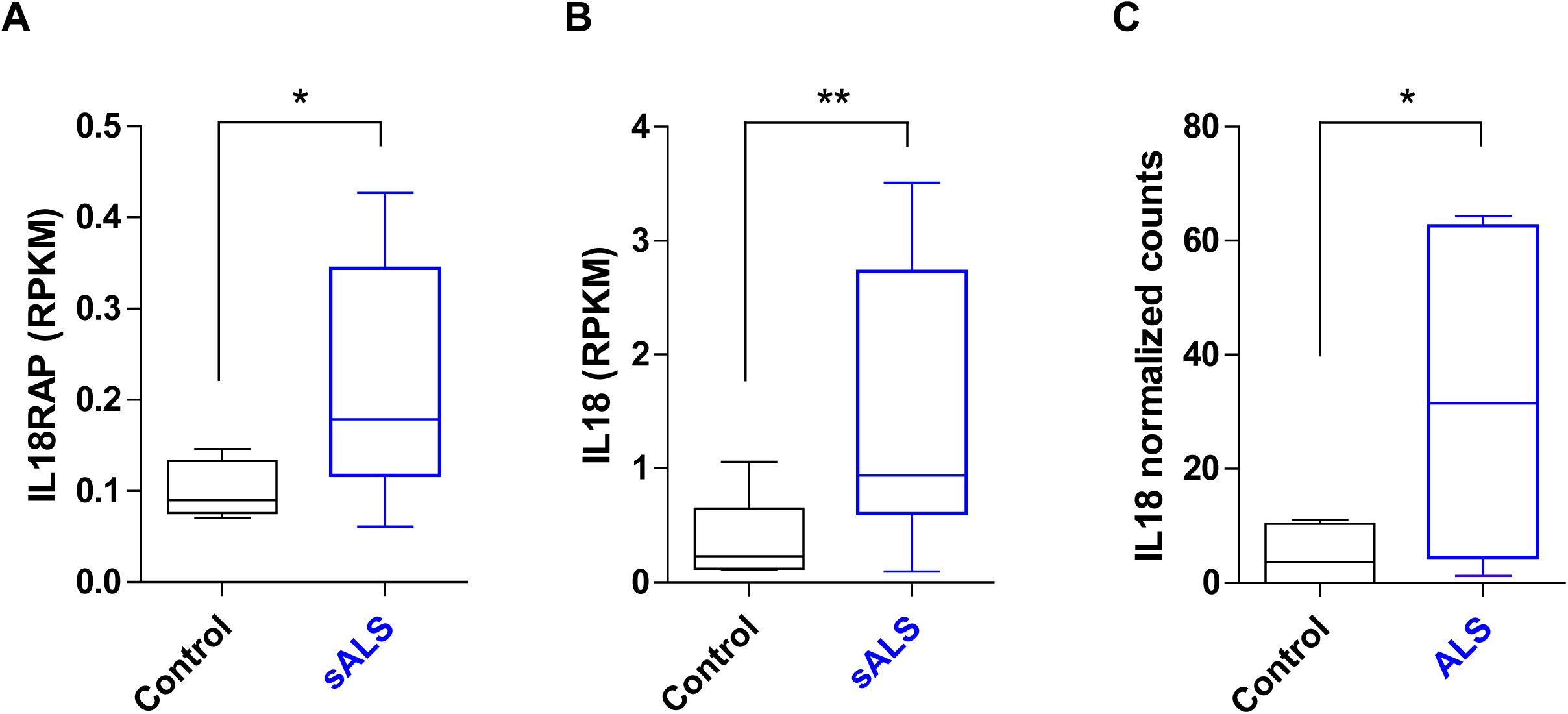
Evaluation of IL18RAP and IL-18 mRNA expression in motor neurons of patients with ALS. **(A-B)** mRNA expression of IL18RAP **(A)** and IL-18 **(B)**, as reads per kilobase million (RPKM), from NGS study of laser capture microdissection–enriched surviving motor neurons from lumbar spinal cords of patients with sALS with rostral onset and caudal progression (n = 12) and non-neurodegeneration controls (n = 9; ^106^ GSE76220). Two-sided Student’s t test with Welch’s correction on log-transformed data. **(C)** IL-18 mRNA expression, as log2- normalized counts, from NGS study of induced ALS motor neurons (n = 4 different donors in duplicates) or non-neurodegeneration controls (n=3 different donors in duplicates; ^107^ DESeq analysis). Box plots depict median, upper and lower quartiles, and extreme points. *P < 0.05; **P < 0.01.

**Supplementary Fig. 9.**
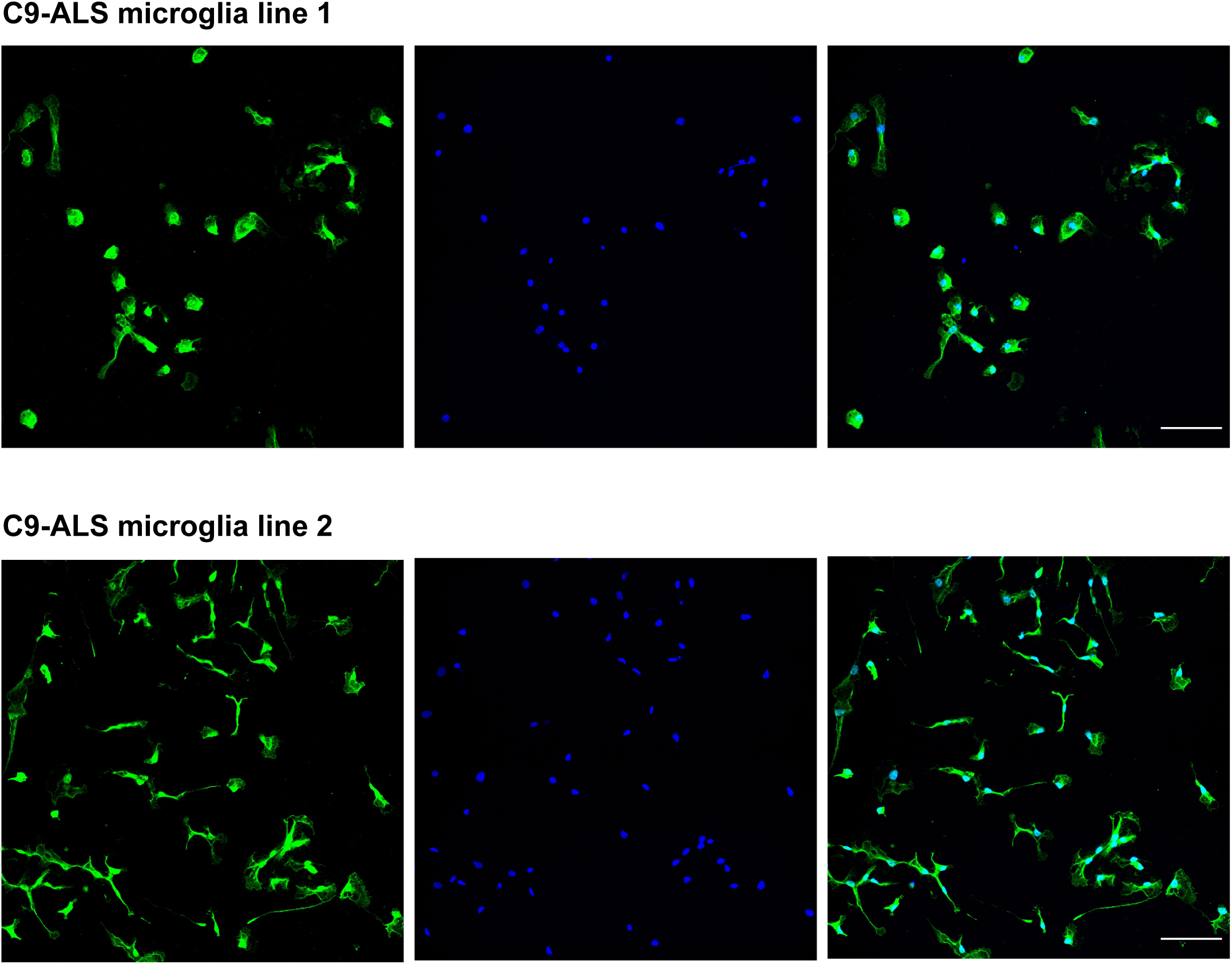
iPSC-derived microglia express the microglial-specific marker, TMEM119. Immunofluorescence staining of TMEM119 (green) and DAPI (blue), in two different C9orf72 iPSC-derived progenitor microglia lines. Lens, ×20; scale bar, 100 μm.

**Supplementary Fig. 10.**
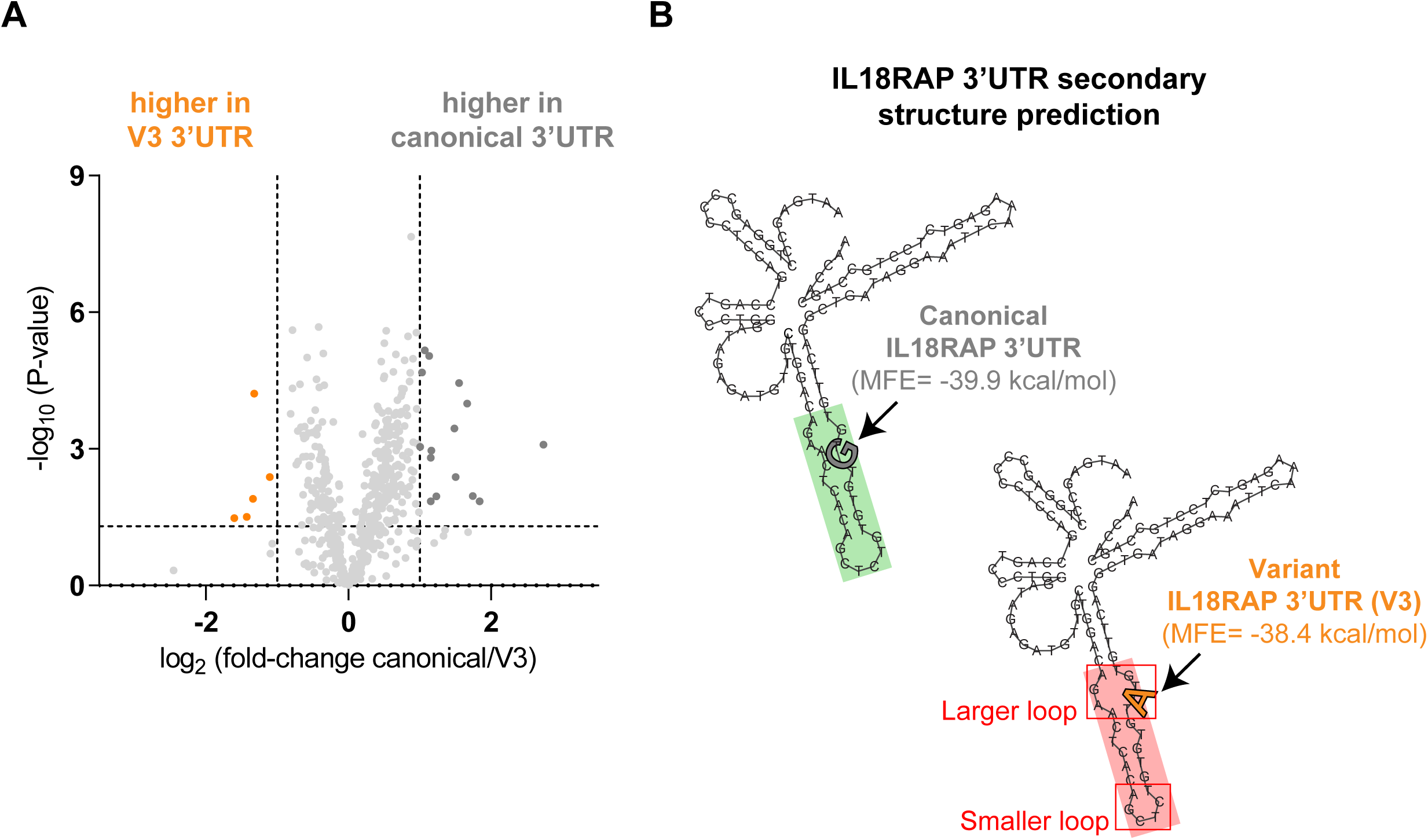
Differentially bound RNA binding proteins to variant 3’UTR (V3) relative to canonical 3’UTR. **(A)** Volcano plot of protein abundance associated with the canonical relative to variant (V3) IL18RAP 3’UTR (x-axis log2 scale), analyzed by MS. Y-axis depicts P-values (−log10 scale). Proteins significantly enriched in association with canonical/variant 3’UTR are colored (grey/orange). Features above the horizontal dashed line demarcate proteins with adjusted p < 0.05, in student’s t-test with FDR correction to multiple hypotheses. Vertical dashed lines are of 2 or ½ fold change (Supplementary Table 9). **(B)** Prediction of 3’UTR secondary structure by RNA Fold ^105^, suggests a minor change to the structure of the sequence harboring a V3 variant (red), relative to the canonical 3’UTR (green).

